# Intrinsic mechanotransduction during apical constriction licenses lineage competence in pluripotent stem cells

**DOI:** 10.1101/2025.11.03.686286

**Authors:** Mehdi Seif Hamouda, Celine Labouesse, George W. Wylde, Daniel Yamamoto, Yekaterina A. Miroshnikova, Srinjan Basu, Clare M. Waterman, Kevin J. Chalut

## Abstract

To acquire the capacity for multi-lineage differentiation, pluripotent stem cells must undergo a transition from naïve pluripotency to lineage competency. This transition requires epithelialization and changes in nuclear architecture. We sought to determine whether the cell and tissue mechanics intrinsic to epithelialization drive pluripotency progression and subsequent lineage commitment to neuroectodermal fate. We demonstrate that naïve mouse embryonic stem cells (mESCs) undergoing early differentiation *in vitro* recapitulate features of epithelialization in the peri-implantation epiblast, specifically apical constriction. We further demonstrate that cell contractility during apical constriction induces a distinct nuclear mechanoresponse, notably enrichment of emerin at the outer nuclear membrane, nuclear envelope localization of SUN2, and the global loss of H3K9me3 heterochromatin which is compensated by H3K27me3. Importantly, these nuclear phenotypes and subsequent neuroectodermal lineage priming require myosin II-mediated contractility, an intact LINC complex, and emerin. We demonstrate that LINC-dependent mechanotransduction through emerin regulates H3K27me3 occupancy on the key early neuroectodermal transcription factor gene, *Sox1*, implicating a mechanical switch in chromatin mediation of neuroectodermal lineage competence. These results indicate that epithelialization-induced nuclear mechanotransduction poises a critical lineage gene for subsequent expression.

**Highlights:**

- Adherent naïve mESCs *in vitro* recapitulate *in utero* apical constriction morphogenesis.
- Contractility, the LINC complex and emerin are required for neuroectodermal lineage competence.
- Contractility and the LINC complex are required for widespread loss of H3K9me3 and gain of H3K27me3 on *Sox1* promoter.

## Introduction

Following fertilization of a totipotent egg, early embryos progress through morulation and blastulation, leading to the emergence of pluripotent cells that form the epiblast of the pre-implantation blastocyst. While pluripotent cells possess the capacity to produce all tissues, they are initially in a so-called naïve state and must undergo a maturation process known as exit from naïve pluripotency to gain lineage competence and sensitivity to lineage-inducing signals (De Belly et al., 2021; Kalkan et al., 2017; Kinoshita et al., 2021; Nichols et al., 2009). This maturation occurs during embryo implantation *in utero,* where naïve pluripotent cells undergo a transition involving a distinct gene expression program and morphogenesis to prime them for lineage commitment. Throughout pluripotency, the transcription factor, *Pou5f1* (OCT4) is expressed, which can facilitate somatic cell reprogramming into induced pluripotent stem cells (Takahashi & Yamanaka, 2006). The expression of other transcription factors that generally function to regulate gene expression networks are associated with naïve pluripotency, such as *Nanog*, and intermediate and primed stages of pluripotency, such as *Otx2* and *Pou3f1* (OCT6) respectively (Acampora et al., 2016; Hackett & Surani, 2014; Kalkan et al., 2017).

In addition to these changes in gene expression profiles, pluripotent cell maturation is also marked by major changes in morphology that include transition from a 3D cell mass in the pre-implantation epiblast to a spherical epithelial monolayer that is marked by cell polarization. This epithelialization morphogenesis requires *Otx2* expression and occurs during implantation of the blastocyst in the uterine wall, where the pluripotent cells of the epiblast initially polarize, perform apical constriction, and form a “rosette” (Shahbazi et al., 2017). Apical constriction of pluripotent cells during implantation is indicated by the central accumulation of phosphorylated non-muscle myosin II light chain in rosette epiblasts of mouse embryos, and can be recapitulated in naïve pluripotent cells during early differentiation in 3D culture (Bedzhov & Zernicka-Goetz, 2014). Subsequently, pluripotent cells undergo lumenogenesis, ultimately forming the epithelial epiblast within the post-implantation blastocyst, also termed the egg cylinder. Pluripotency concludes at the onset of gastrulation, where lineage commitment to the germ layers begins (Shparberg et al., 2019). Although the genetic and morphological changes that occur during early embryogenesis are well-described, whether they are interdependent is not known.

The nucleus has become increasingly appreciated in its role in converting mechanical stimuli experienced by cells during morphogenesis into changes in molecular interactions and gene expression, in a process called nuclear mechanotransduction. This process is mediated by either extrinsic forces on cells or intrinsic cytoskeletal contractility, with force transmission to the nucleus, in part, mediated by the Linker of the Nucleoskeleton and Cytoskeleton (LINC) complex (Arsenovic et al., 2016; Crisp et al., 2006; Haque et al., 2006; Padmakumar et al., 2005). The LINC complex is composed of cytoskeleton-binding KASH (Klarsicht, ANC-1, Syne Homology) domain-containing proteins called nesprins (Nuclear Envelope Spectrin Repeat-containing Proteins) that reside in the outer nuclear membrane (ONM), which also bind in the space between the ONM and the inner nuclear membrane (INM) to the SUN (Sad1/UNC-84) domain of trans-INM SUN proteins. Mechanical stimuli impinged on LINC can enable forces transmitted from the cytoskeleton through the nuclear envelope (NE) to drive force-induced changes in nuclear architecture by influencing nuclear proteins, such as nuclear lamins, histone modifying enzymes, and chromatin organization (Hervé & Miroshnikova, 2024; Miroshnikova & Wickstrom, 2022; Nava et al., 2020; Stephens & Miroshnikova, 2024; Tajik et al., 2016). Emerin, also important to nuclear mechanotransduction, typically resides in the INM where it can bind directly to the luminal domain of SUN proteins (Fernandez et al., 2022; Guilluy et al., 2014; Lammerding et al., 2005). Emerin can also recruit Polycomb Repressive Complex 2 (PRC2) component EZH2 and HDAC3 to the nuclear periphery where they can promote histone modifications that regulate gene accessibility (Demmerle et al., 2012, 2013; Haque et al., 2010). There is emerging evidence that nuclear mechanotransduction can regulate gene expression and cell fate during later stages of development and differentiation (Biedzinski et al., 2020; Carley et al., 2021; Discher et al., 2007; Le et al., 2016; Poleshko et al., 2017; Shiraishi et al., 2023). While numerous studies have demonstrated functional consequences of cells interacting with extracellular matrices of differing physical parameters or subjected to externally applied forces (Han et al., 2020; Labouesse et al., 2021; Le et al., 2016; Nava et al., 2020; Verstreken et al., 2019), it remains unclear whether intrinsic, cell-generated forces during early differentiation can also elicit nuclear responses that control gene expression. Whether nuclear mechanotransduction plays a role in regulating gene expression during the morphogenic events of early embryogenesis has not been explored.

There are hints of evidence that nuclear mechanotransduction may play a role in the transition of stem cells from naïve pluripotency to lineage committed in early embryogenesis, but it is far from conclusive. For example, pluripotent cells in the naïve state are largely insensitive to applied stretch, only showing mechanically-regulated gene expression responsiveness in later stages of pluripotency, suggesting a potential role for mechanical regulation in lineage commitment (Verstreken et al., 2019). Coincident with the onset of mechanoresponsiveness, differentiating pluripotent cells change their nuclear mechanics which is tied to widespread changes in chromatin compaction (Chalut et al., 2012; Pagliara et al., 2014). Chromatin changes could be attributed to alterations of constitutive heterochromatin-associated histone post-translational modifications, such as repressive H3K9me3, and facultative heterochromatin marks, such as H3K27me3, which plays an essential role in the regulation of gene expression in early development (Boyer et al., 2006; Padeken et al., 2022; Saksouk, Simboeck, et al., 2015; Trojer & Reinberg, 2007; Wiles & Selker, 2017). Interestingly, pluripotent cells of peri-implantation mouse embryo explants show a transient period of widespread H3K9me3 loss compensated by H3K27me3, which is a phenotype similar to mechanically stimulated cells, and consistent with early observations of pluripotent cells with depleted H3K9me3 (Le et al., 2016; Nava et al., 2020; Neagu et al., 2020; Peters et al., 2003; Saksouk, Barth, et al., 2015; Song et al., 2022; Xu et al., 2023). These findings culminate into compelling questions as to whether these nuclear architecture changes, reminiscent of mechanotransduction phenotypes, are driven by morphogenesis.

Supported by this background, we hypothesized that morphogenesis during early differentiation in pluripotent cells are linked with changes in the nucleus critical for pluripotency maturation and early lineage commitment. To directly investigate the relationship between the cell-generated forces during epithelialization and nuclear mechanotransduction, we studied actin cytoskeleton remodeling during early differentiation of naïve pluripotent mouse embryonic stem cells (mESCs) in an *in vitro* model system. Here we show that adherent mESCs *in vitro* recapitulate elements of apical constriction that occur in the epiblast during embryo implantation. We find that the actomyosin contractility that occurs during apical constriction drives changes in nuclear envelope proteins and histone post-translational modifications, featuring changes in SUN2 and emerin localization and a switch between H3K9me3 and H3K27me3 histone marks. These changes, which are mediated by the LINC complex, are coupled to the regulation of neuroectodermal lineage master transcription factor, *Sox1*, licensing it for later expression. Our study uncovers a potential role of tissue morphogenesis-driven nuclear mechanotransduction pathways that enable regulation of lineage gene expression, carrying implications for developmental processes that involve distinct morphogenetic processes with subsequent fate choices.

## Results

### Adherent mESCs Recapitulate Apical Constriction Morphogenesis of the Peri-implantation Epiblast During Naïve Pluripotency Transition

We first sought to establish an adherent cell model to allow high-resolution microscopy during morphogenesis in pluripotent cells. We utilized mESCs, which can model early pluripotency differentiation by the withdrawal of “2i” (MEK/ERK and GSK3) inhibitors from a chemically-defined culture system, closely recapitulating the changing transcriptional profiles of developing *in utero* counterparts, while featuring a default commitment to the neuroectodermal lineage (Argelaguet et al., 2019; Mulas et al., 2019; Yang et al., 2019; Ying et al., 2008). Furthermore, differentiating mESCs in 3D suspended 2i culture recapitulate features of epithelialization morphogenesis, characterized by rosette formation of cells with contractile actin enrichment in the center (Bedzhov & Zernicka-Goetz, 2014; Christodoulou et al., 2018; Kim et al., 2022; Mole et al., 2021; Neagu et al., 2020; Shahbazi et al., 2017). To determine if adherent mESCs undergo actomyosin contractility similar to that *in utero* and in 3D culture, we conducted time-course experiments and collected protein lysates from mESCs after 2i withdrawal to induce exit from naïve pluripotency and early differentiation (Figure S1A). As expected, at around ∼12 hours into differentiation, expression of the intermediate pluripotency transition marker, OTX2, was upregulated (Figure 1A) (Kalkan et al., 2017; Kinoshita et al., 2021; Neagu et al., 2020). The initial upregulation of OTX2 was preceded by a rise in myosin II light chain phosphorylated on serine 16 (PMLC2), a marker of non-muscle myosin II activity and a proxy for cellular contractility (Vicente-Manzanares et al., 2009). Myosin II contractility peaked and began to decline before the peak of OTX2 expression was reached. This indicated that cells in early differentiation transition undergo a transient window of increased actomyosin contractility that precedes a rise in OTX2 expression, suggesting that contractility could promote this driver of peri-implantation epiblast morphogenesis (Shahbazi et al., 2018).

**Figure 1.**
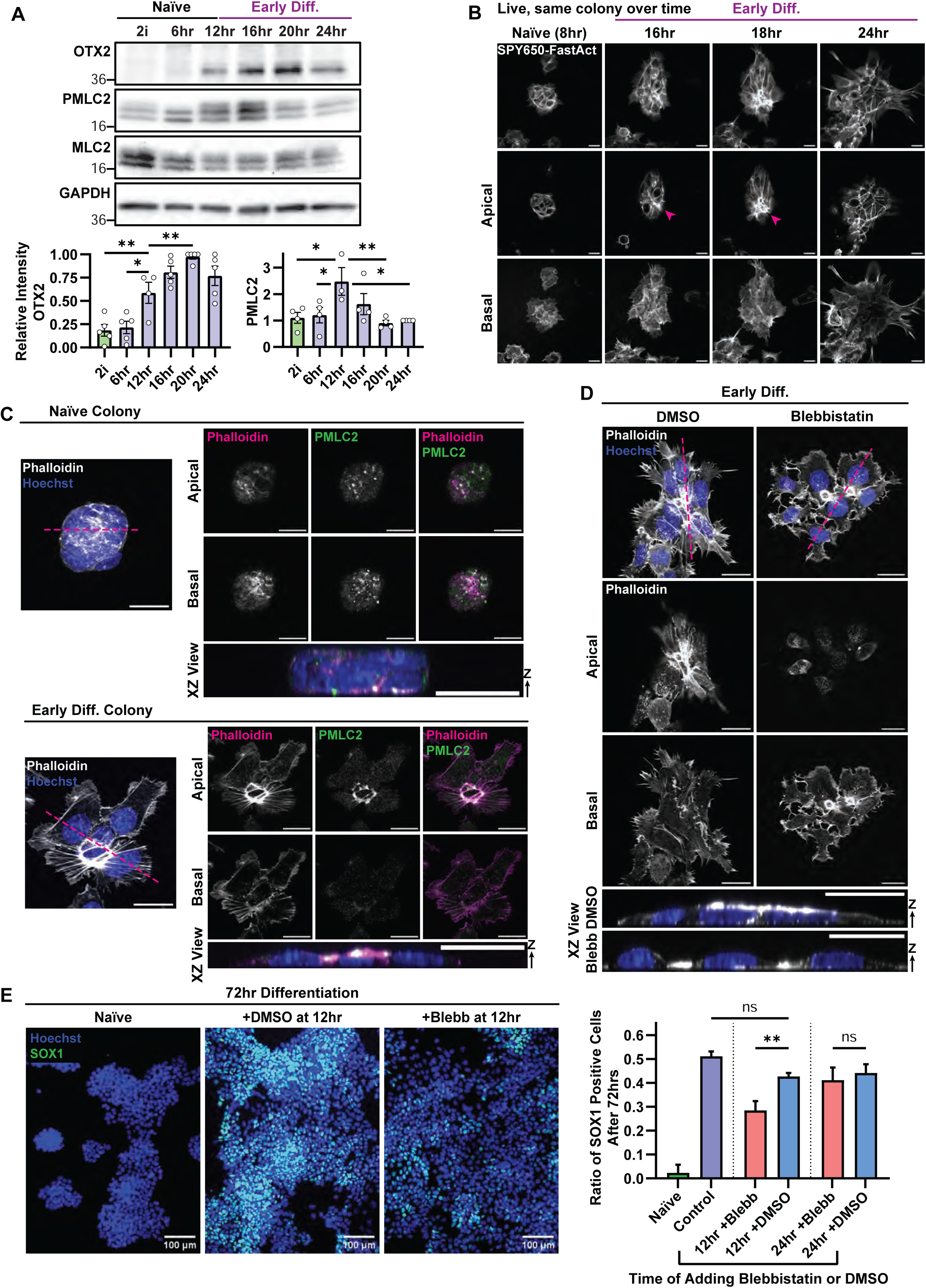
Apical Constriction morphogenesis is recapitulated by adherent mESCs during the exit from naïve pluripotency and contractility is required for efficient differentiation to neuroectoderm. **(A)** Above: Representative western blot of lysates of mESC cultured in media containing inhibitors (2i) to maintain naïve pluripotency (Naïve) or mESC collected at noted timepoints hours after 2i withdrawal (6, 12, 16, 20 and 24 hours) to allow exit from naïve pluripotency and initiation of early differentiation (Early Diff). Blots were probed for OTX2, myosin II light chain phosphorylated on serine 16 (PMLC2), myosin II light chain (MLC2) and GAPDH. Below: Quant i ficat i on of blots like those presented above. Lef t : Normalized to GAPDH and relative max level of OTX2. Right: Normalized to MLC2 and relative to level of PMLC2 at 24 hr. (mean ± SEM of N=4 independent experiments, Ordinary one-way ANOVA, Dunnett Multiple Comparisons Test, OTX2: **P=0.0059, *P=0.0115, **P=0.0083, PMLC2: *P=0.0184, *P=0.0300, **P=0.0068, *P=0.0110). **(B)** Images from a time-lapse 3D confocal movie of live, adherent, mESC colonies at the noted times after 2i withdrawal to allow exit from naïve pluripotency and initiation of early differentiation (Naïve, Early Diff) with F-actin labelled with SPY650-FastAct (greyscale). Top rows show 3D projections of confocal Z-stacks, second and third rows show apical and basal confocal images, respectively. Pink arrowheads point to bright apical F-actin ring structures. Scale bars, 10μm. **(C, D)** mESC colonies fixed in 2i (Naïve) or 16 hours after 2i withdrawal (Early Diff) to allow exit from naïve pluripotency and initiation of early differentiation and stained with fluorescently-tagged phalloidin to visualize F-actin (grayscale or magenta in **C,** grayscale in **D**) and Hoechst to visualize DNA (blue) with PMLC2 additionally immunostained in (**C,** green). Pink dashed lines highlight the plane of the XZ view from the 3D reconstruction in the bottom row. In **(D),** colonies were treated at 12 hours with 2μM Blebbistatin or DMSO vehicle control. Left panels in **(C)** and top row in **(D)** show projections of 3D confocal Z-stacks. Top two rows in **(C)** and second and third rows in **(D)** show apical and basal confocal images, respectively. Scale bars, 20μm. **(E)** Left three panels: Epifluorescence images of mESCs treated with 2μM blebbistatin or DMSO vehicle at 12 hours after 2i withdrawal to allow exit from naïve pluripotency and initiation of early differentiation, or kept in 2i (Naïve), then fixed after 72 hours, with Hoechst-stained DNA (blue) and immunostained for SOX1 (green). Scale bar, 100μm. Right panel: SOX1 expression was quantified as ratio of SOX1 positive cells. (mean ± SD of N=3 independent experiments, ordinary one-way ANOVA with Tukey multiple comparisons test, \*\**P*=0.002).

Next, to determine if increased actomyosin contractility in early differentiation actually coincided with cell morphogenesis, we performed time-lapse 3D confocal imaging of the actin cytoskeleton, with SPY650-FastAct labeled filamentous actin (F-actin), in living adherent mESCs during early differentiation after 2i withdrawal (Figure 1B, Video S1). In naïve or at very early differentiation timepoints, cell colonies presented typical dome shapes with F-actin delineating individual cells and concentrated in a band around the periphery of the colony at the base, and within basal filopodia. Starting around 16 hours after 2i removal, colonies notably formed small (∼2-8µm), dense ring-like F-actin rich structures on the apical side in colony centers. We confirmed formation of apical ring-like F-actin rich structures in fixed colonies with fluorescent phalloidin-labeled F-actin (Figure S1B). The apical F-actin rings often featured emanating radial stress fibers that terminated at cell adhesions on the basal side at the colony periphery. These apical F-actin rings appeared to be formed from the converging apical surfaces of multiple adjacent individual cells that were joined by a thick F-actin bundle that spanned between cells, reminiscent of the circumferential ’purse string’ model of apical constriction (Martin & Goldstein, 2014). At later timepoints, the apical surfaces expanded, and colonies resembled the apical surfaces of epithelial cells (Figure 1B, Video S1). Thus, exit from naïve pluripotency is accompanied by drastic remodeling of the actin cytoskeleton marked by an apical ring-like structure.

To determine if increased myosin II activity and appearance of apical F-actin rings were associated with apical constriction, we immunostained for PMLC2 in F-actin-labelled colonies in cells fixed after removal of 2i (Figure 1C). Along with colonies flattening during early differentiation, 3D confocal imaging showed that the apical F-actin rings present in colonies during early differentiation strongly colocalized with PMLC2. We also observed PMLC2-rich, apical F-actin structures in colonies as small as two cells, further suggesting this is a cell-intrinsic occurrence during early differentiation (Figure S1C). Similarly, immunostaining for non-muscle myosin II heavy chain A and B isoforms (MHCIIA, MHCIIB) in early differentiating mESCs also revealed strong colocalization with apical, F-actin rich structures, along with basal F-actin stress fibers and lamellae (Figure S1D, S1E). Preferential enrichment of PMLC2 in apical actin structures indicated the polarized targeting and activation of non-muscle myosin II to drive contractility. These results showed that adherent mESCs undergoing an early differentiation transition recapitulate aspects of apical constriction morphogenesis that occur in the peri-implantation epiblast.

### Morphogenesis-Associated Apical Constriction Enables Efficient Neuroectodermal Lineage Commitment

We next tested for the functional impact of mESC colony apical constriction on changes in gene expression. To accomplish this, we investigated the role of contractility during early differentiation and subsequent neuroectodermal lineage priming. Treatment of mESC colonies with blebbistatin to block non-muscle myosin II ATPase activity after 12 hours relative to 2i removal, even at a concentration low enough to not impact proliferation (2μM), resulted in flattened colonies, inhibited cell-cell contacts, drove formation of peripheral lamellipodia, and eliminated the appearance of apical constriction features at later timepoints (Figure 1D, S1F). Interestingly, however, multicellular F-actin ring structures still formed, although they were located at the basal cell plane. In contrast, apical constriction and radial stress fibers were maintained in colonies treated with DMSO vehicle control. This indicated that actomyosin contractility is critical for maintenance of cell-cell adhesion and apical constriction during exit from naïve pluripotency.

Due to the temporal overlap of mESC colony apical constriction with expression of the early differentiation marker OTX2, we next asked if actomyosin contractility was required for the initial exit from the naïve pluripotent state. We probed for four important factors that regulate and track the stages of exit from pluripotency, *Nanog*, *Otx2*, *Pou5f1* (OCT4), and *Pou3f1* (OCT6), using an RT-qPCR time-course after removal of 2i in the presence of DMSO or 2μM blebbistatin (Figure S1G). In all cases, the expression level and timing of these genes marking pluripotency progression were unaffected by blebbistatin treatment. This indicated that exit from naïve pluripotency, and entry into the following intermediate pluripotency phase (the ‘formative phase’) does not depend on actomyosin contractility and mESC colony apical constriction.

Given that myosin II inhibition had no effect on exit from naïve pluripotency and maturation of mESCs, we next asked if apical constriction-associated contractility was instead required for efficient transition to the next stage: neuroectodermal lineage priming. We thus measured expression of master neuroectodermal lineage transcription factor and earliest lineage marker, SOX1, and analyzed the effects of inhibiting contractility. Blebbistatin treatment at 2μM, 12 hours relative to 2i removal, resulted in a significantly reduced percentage of SOX1-positive cells at a much later time-point (72 hours) compared to DMSO vehicle control (Figure 1E). Intriguingly, when actomyosin contractility was inhibited after the period of apical constriction (after 24 hours relative to 2i removal), SOX1 expression at 72 hours proceeded similarly to the control conditions. These results suggest that there is a time window during early differentiation between 12 and 24 hours, coinciding with apical constriction, where contractility is required for efficient SOX1 expression at a significantly later time-point (72 hours).

### Apical Constriction Is Associated with Peri-nuclear Actin and Nuclear Deformations

Given how stresses in the actin cytoskeleton have been implicated in nuclear mechanotransduction (Le et al., 2016; Miroshnikova & Wickstrom, 2022), we next asked if apical constriction during early differentiation could directly mechanically impinge on nuclei. 3D confocal analysis of naive and differentiating colonies of mESCs showed that apical actomyosin structures in early differentiating colonies caused large-scale deformations on the apical surfaces of underlying nuclei (Figure 2A). These deformations coincided with apical F-actin rings as well as taut radial stress fibers. Furthermore, we observed F-actin networks in contact with the nuclear surface in naïve mESCs, and quantification showed that these significantly increased in density during early differentiation, coinciding with apical constriction, with some nuclei becoming completely and tightly enveloped in an F-actin shroud (Figure 2B, C). Additionally, inhibition of actomyosin contractility with blebbistatin eliminated apical nuclear deformations and the association of F-actin networks with the surface of nuclei. These results show that apical constriction is associated with the de-novo formation of a peri-nuclear F-actin shroud and tensed F-actin cables that directly induce nuclear deformations.

**Figure 2.**
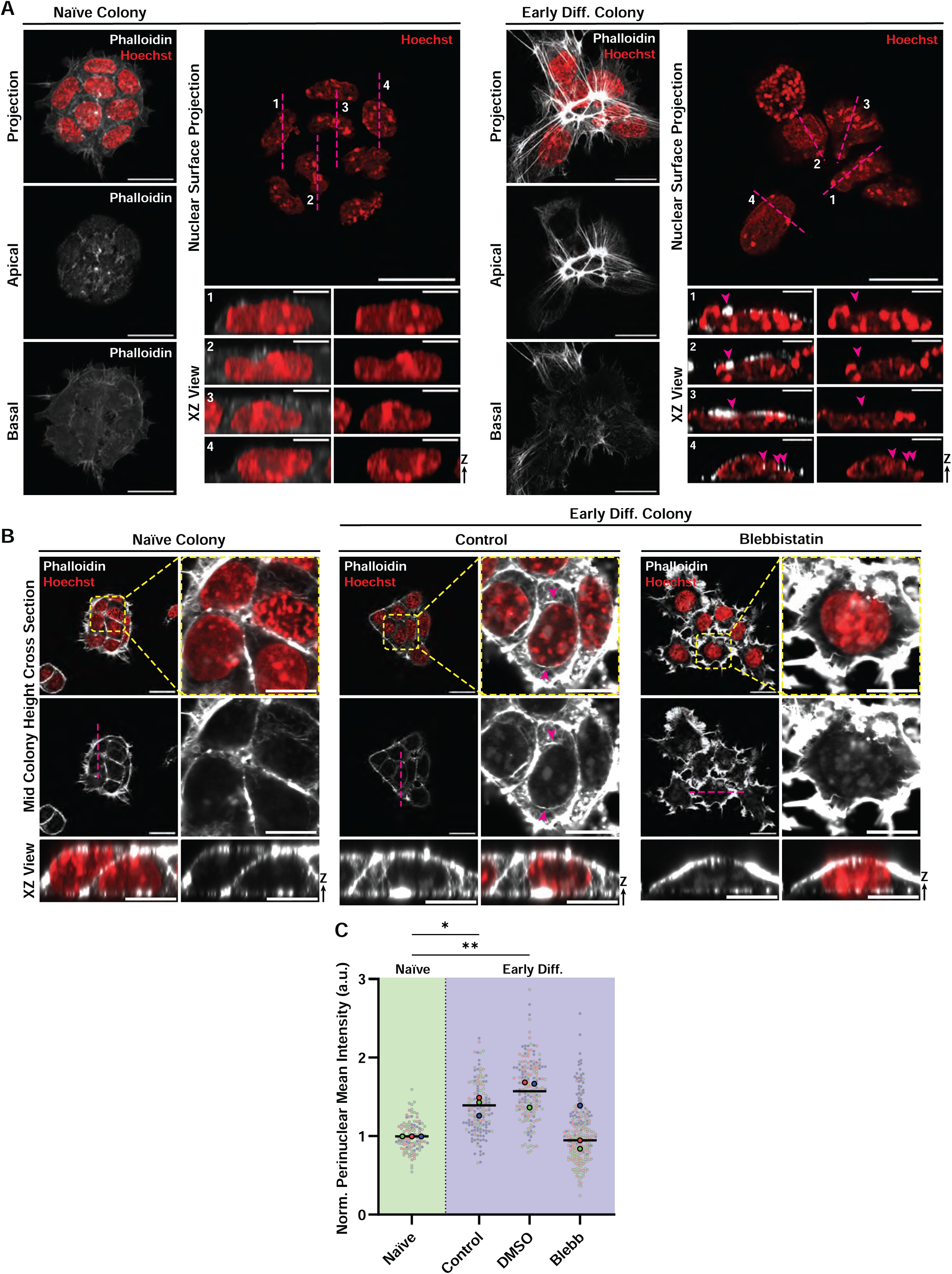
Adherent colony apical constriction is associated with F-actin induced nuclear deformation and the formation of perinuclear actin networks. **(A, B)** Super-resolution confocal imaging of fixed mESCs cultured in media containing inhibitors (2i) to maintain naïve pluripotency (Naïve) or mESC at ∼16 hours after 2i withdrawal to allow exit from naïve pluripotency and initiation of early differentiation (Early Diff) stained with fluorescently-tagged phalloidin to visualize F-actin (grayscale) and Hoechst to visualize DNA (red). **(A)** Upper left panels show projections of 3D confocal Z-stacks. Middle left and bottom left show apical and basal confocal images, respectively. Upper right shows a projection of a surface rendering from a 3D confocal stack. Bottom right four rows show numbered XZ views from a 3D reconstruction, taken along the numbered, pink dashed lines above. Pink arrows indicate apical actin structures colocalizing with deformations of underlying apical nuclear surfaces. Scale bars, 20μm, zoomed XZ view scale bars, 5μm. **(B)** Upper two rows: Confocal images taken at mid-colony height to visualize perinuclear f-actin in the presence of 2μM blebbistatin or DMSO vehicle (Control) added after 12 hours. Yellow dashed boxed regions in upper right panels shown zoomed at left. Pink dashed lines highlight the plane of the XZ view from 3D reconstructions in the bottom row. Pink arrows indicate regions of enriched f-actin on the nuclear surface. **(C)** Quantification of perinuclear actin intensity from 4-pixel wide line scans taken around the NE perimeter in the central confocal planes **(B)**, (mean ± SD of N=3 independent experiments, colored points correspond to individual cells within each experiment, Ordinary one-way ANOVA with Tukey multiple comparisons test, \**P*=0.0488, \*\**P*=0.0071).

### Apical Constriction Associated Contractility Enhances Neuroectodermal Lineage Commitment via the LINC complex

We next sought to determine how contractile stresses generated in the cytoskeleton are transmitted to the nucleus to regulate neuroectoderm lineage commitment during early differentiation. We thus examined the contributions of NE proteins known to mediate nuclear mechanotransduction, including the LINC complex and emerin, in this process. To perturb the LINC complex, we produced a cell line that allowed doxycycline-inducible expression of a truncated nesprin-2 mutant that lacks an actin binding domain, called DN-SRKASH, that decouples the LINC complex in a dominant negative manner by outcompeting endogenous nesprins for binding to SUN proteins (Lombardi et al., 2011; Luxton et al., 2010). Indeed, expression of DN-SRKASH caused loss of the LINC component SUN2 and the perinuclear f-actin meshwork from the NE (Figure 3A, S2A), in agreement with nesprin proteins’ known role in formation and stabilization of trans-NE LINC complexes (Crisp et al., 2006; Padmakumar et al., 2005). Heterogeneity of DN-SRKASH expression revealed a significant negative correlation between DN-SRKASH expression and SUN2 levels at the NE (Figure S2B). Similarly, inhibition of actomyosin contractility with blebbistatin also reduced levels of SUN2 at the NE in early differentiating colonies (Figure 3B), supporting the contention that contractility is required for stabilizing LINC in the NE (Niu et al., 2022). However, unlike blebbistatin treatment, DN-SRKASH expression did not inhibit colony formation and apical constriction during early differentiation, with apical F-actin structures still displaying distinct PMLC2 enrichment (Figure S2C), although perinuclear actin did not significantly change compared to naïve condition (Figure S2D). These results demonstrate the specificity of our perturbation and further show that apical constriction is upstream of formation and stabilization of the LINC complex in the NE through recruitment of SUN2.

**Figure 3.**
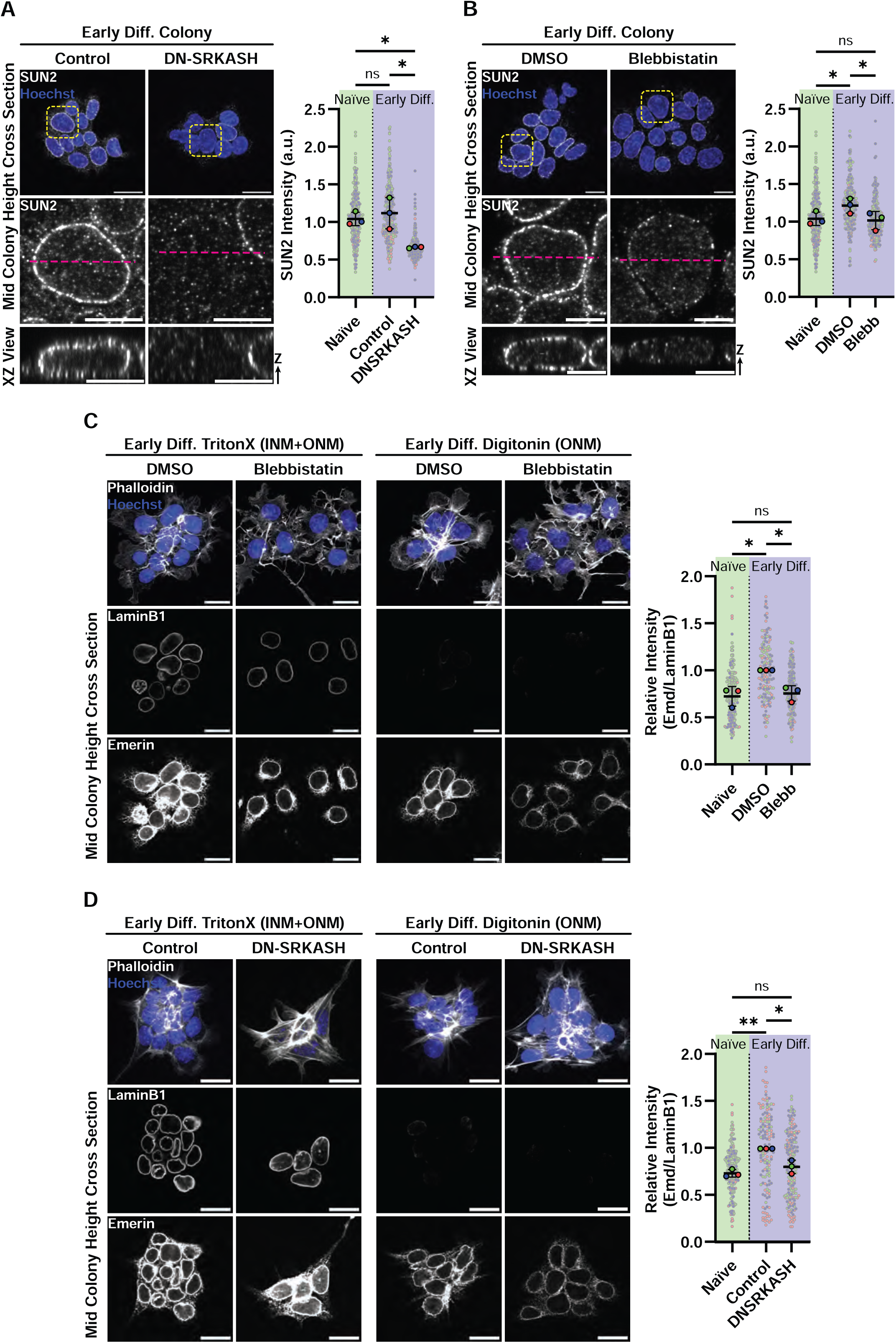
Actomyosin contractility and the LINC complex are required for SUN2 NE localization and enrichment of ONM emerin during early differentiation. **(A, B)** Left panels: Confocal images taken at mid-colony height of mESCs fixed at 16 hours after 2i withdrawal to allow exit from naïve pluripotency and initiation of early differentiation (Early Diff) immunostained for SUN2 (grayscale) and stained with Hoechst to visualize DNA (blue) either **(A)** of cells lacking (control) or bearing inducible RFP-tagged DN-SRKASH with doxycycline treatment, or **(B)** in the presence of 2μM blebbistatin or DMSO vehicle control. Top row: yellow dashed boxes denote regions zoomed in the second row, bottom rows show XZ view of a 3D reconstruction. Pink dashed lines highlight the plane of the XZ view of the 3D images in the bottom row. Right panels: Quantification of mean intensity of SUN2 from max intensity Z projections of individual nuclei. (mean ± SD, N=3 independent experiments, colored points correspond to individual cells within each experiment, Repeated Measures One-Way ANOVA, Tukey Multiple Comparisons Test, Blebbistatin: *P=0.0470, *P=0.0313, DNSRKASH: *P=0.0304, *P=0.0161). **(C, D)** Confocal images of mESCs fixed at 16 hours after 2i withdrawal to allow exit from naïve pluripotency and initiation of early differentiation (Early Diff) immunostained for emerin (bottom row) and LaminB1 (middle row), (both taken at mid-colony height), and also stained with fluorescent phalloidin to label F-actin (grayscale, top row) and Hoechst to visualize DNA (blue), (top row, 3D projection of confocal stack) either **(C)** in the presence of 2μM blebbistatin or DMSO vehicle control or **(D)** of cells lacking (control) or bearing inducible RFP-tagged DN-SRKASH with doxycycline treatment. Cells were permeabilized with tritonX-100 (left two columns) to allow antibody labelling of both the INM and ONM, or digitonin (middle two columns) to allow antibody labelling of the ONM but not the INM. Right panels: Quantification of outer nuclear membrane emerin in images from digitonin digest conditions, normalized (relative to control) mean intensity ratio of emerin to LaminB1, of max intensity Z projections of individual nuclei (mean ± SD, N=3 independent experiments, colored points correspond to individual cells within each experiment, Repeated Measures One-Way ANOVA, Tukey Multiple Comparisons Test, Blebbistatin: *P=0.0324, *P=0.0474, DNSRKASH: **P=0.0062, *P=0.0186), Scale bars, 20μm.

The LINC complex can interact with emerin to elicit nuclear mechanoresponses, causing its relocation from the INM to the ONM (Essawy et al., 2019; Guilluy et al., 2014; Le et al., 2016). To determine if apical constriction regulates emerin localization, we employed a selective permeabilization protocol to allow immunostaining for emerin in either only the ONM or in both the INM and ONM (Adam et al., 1990; Haque et al., 2010). This showed that emerin resided primarily in the INM in naïve mESCs but became enriched in the ONM in colonies with features of apical constriction during early differentiation (Figure 3C, S2E). This ONM emerin enrichment required contractility and the LINC complex, as enrichment was reduced by blebbistatin treatment or DN-SRKASH expression (Figure 3C, 3D). Together, our results indicate that apical constriction-associated contractility during early differentiation in mESCs enhances formation of the LINC complex, which in turn drives enrichment of emerin in the ONM and assembly of an F-actin network on the nuclear surface.

### Apical Constriction Associated Contractility Enhances Neuroectodermal Lineage Commitment via the LINC complex

We next sought to determine whether early lineage commitment could be mediated by LINC- and emerin-dependent nuclear mechanotransduction. We first checked whether the LINC complex regulated pluripotency progression. We found, similar to the effects of inhibiting contractility, that perturbation of the LINC complex with DN-SRKASH overexpression at the time of 2i withdrawal had no impact on expression of Nanog, Otx2, *Pou5f1* (OCT4), or *Pou3f1* (OCT6), factors that regulate pluripotency maturation (Figure S2F, S2G). In contrast, we found that DN-SRKASH expression induced at 0 or 12 hours after 2i removal resulted in impaired SOX1 expression after 72 hours (Figure 4A). To ensure that this was caused by defects in the LINC complex specifically, we also generated a cell line with an inducible, dominant-negative SUN1 mutant that lacks a transmembrane domain (DN-SUN1) and thus decouples the LINC complex similar to DN-SRKASH. DN-SUN1 expression at early timepoints also gave an impaired SOX1 expression phenotype when expressed prior to or during early differentiation (Figure 4B) (Hennen et al., 2018). Interestingly, inducible overexpression of *Lmna* (corresponding to LAMIN A/C), which is normally at very low levels in mESCs and other pluripotent cells, also resulted in an impaired SOX1 phenotype when expressed at up to 12hr after 2i removal (Figure S3A) (Constantinescu et al., 2006; Eckersley-Maslin et al., 2013). Since contractility and LINC perturbations caused changes in emerin localization in the NE (Figure 3C, D), we also tested the requirement for emerin in early differentiation and neuroectoderm commitment. Confocal imaging in mESCs transfected with emerin siRNA showed normal apical constriction morphogenesis and RT-qPCR of pluripotency progression factors (*Nanog* and *Otx2*) showed normal gene expression timing and levels during early differentiation (Figure S3B, S3C). On the other hand, knockdown of emerin with two different siRNAs inhibited neuroectodermal lineage priming, as indicated by reduced expression of SOX1 at 72 hours after 2i removal (Figure 4C), while non-targeting controls (NTC) had no effect. Together, these results suggest that there is a time window during early differentiation coincident with apical constriction during which priming of *Sox1* is mediated by LINC-, emerin- and contractility-mediated nuclear mechanotransduction for expression later in development to mediate neuroectodermal lineage commitment.

**Figure 4.**
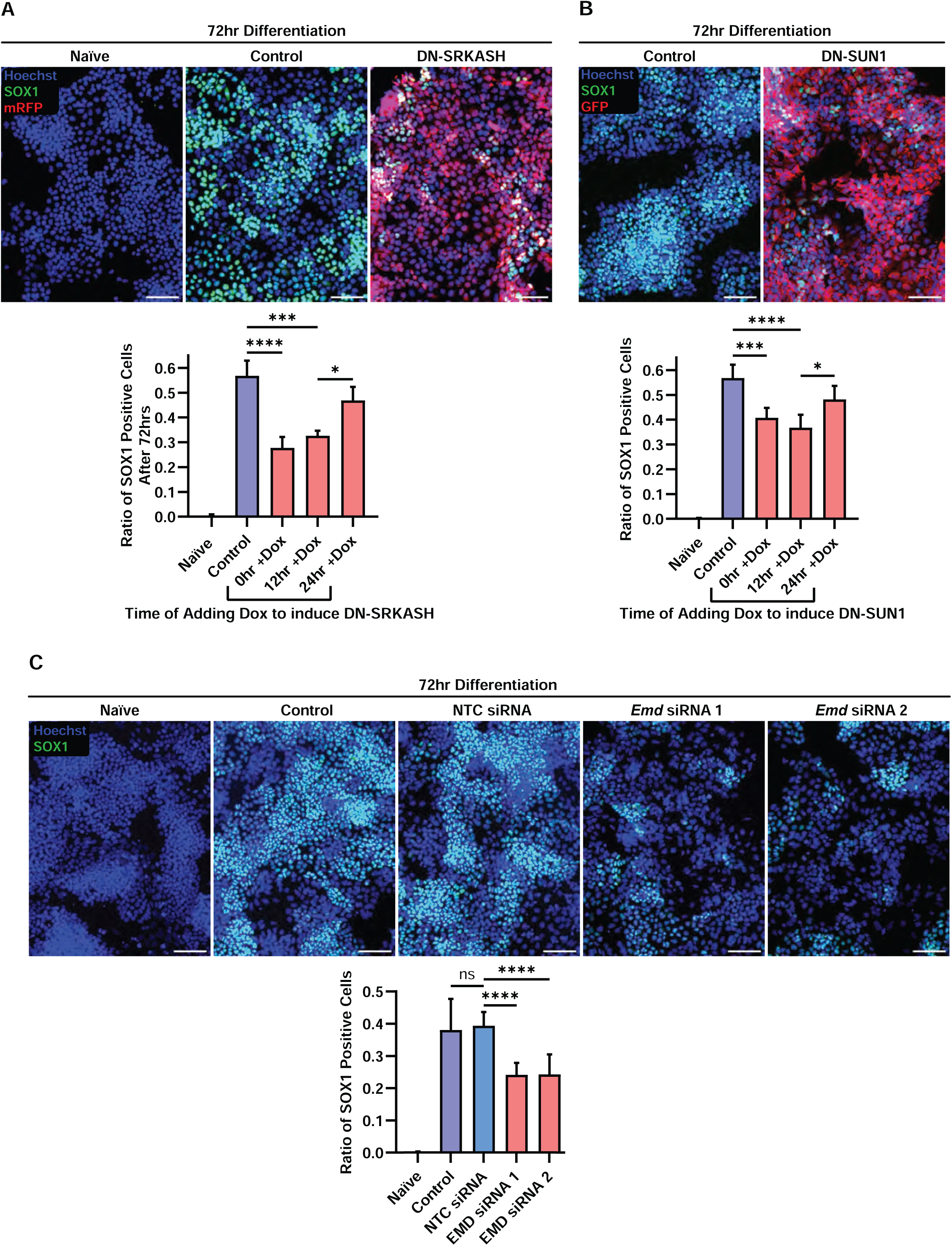
The LINC complex and emerin are required during an early differentiation window for efficient differentiation to neuroectoderm. **(A,B,C)** Top rows: Epifluorescence images of mESCs fixed after 72 hours that were cultured in media containing inhibitors (2i) to maintain naïve pluripotency (Naïve) or after 2i withdrawal to allow exit from naïve pluripotency and early lineage commitment, immunostained for SOX1 (green) and stained with Hoechst to visualize DNA (blue), for cells lacking (control) or bearing doxycycline (Dox)-inducible RFP-tagged DN-SRKASH (red, **A**) or GFP-tagged DN-SUN1 (red, **B**) or mock-transfected (control) or transfected with either non-targeting siRNAs (NTC) or siRNAs targeting two different sequences within emerin (Emd siRNA 1 and Emd siRNA 2**, C**). Scale bars, 100μm. Bottom rows: Quantification of the fraction of SOX1-positive cells fixed after 72 hours in 2i inhibitor (Naïve) or after 2i withdrawal to allow exit from naïve pluripotency and initiation of early differentiation in cells like those described above. **(A)** mean ± SD of N=3 independent experiments, Ordinary one-way ANOVA with Tukey multiple comparisons test, ****P<0.0001, ***P=0.0003, *P=0.015; **(B)** mean ± SD of N=5 independent experiments, Ordinary one-way ANOVA with Tukey multiple comparisons test, ***P<0.0003, ****P=<0.0001, *P=0.0185; **(C)** mean ± SD of N=3 independent experiments, Ordinary one-way ANOVA with Tukey multiple comparisons test, ****P=<0.0001, ****P=<0.0001).

### Apical Constriction Influences Histone Post-Translational Modifications Via Actomyosin Contractility And The LINC Complex

Given our finding that apical constriction and NE-associated proteins regulate lineage commitment during early differentiation, together with the well-supported notion that mechanical stimulation of nuclei promotes histone post-translational modifications to modulate gene expression in other systems (Le et al., 2016; Nava et al., 2020; Song et al., 2022; Xu et al., 2023), we next asked if there is a link between apical constriction, nuclear mechanotransduction, and heterochromatin regulation in mESCs during early differentiation. We first tested the role of apical constriction in the level of the strongly repressive modification, H3K9me3, the reduction of which could prime chromatin for subsequent gene expression during lineage commitment. Immunostaining of naive mESCs for H3K9me3 together with Hoechst staining of DNA and phalloidin staining of F-actin revealed an expected enrichment of this repressive modification at the nuclear periphery and in Hoechst-rich chromocenters in the nuclear interior in all cells within colonies (Figure 5A, S4A, S4B). Initiation of exit from naive pluripotency and early differentiation was marked by notable cell-to-cell variability in H3K9me3 intensity depending on position within the colony, with cells in the interior of colonies displaying lower H3K9me3 compared with cells on the colony periphery (Figure 5A). Indeed, comparing H3K9me3 levels with F-actin staining showed that nuclei in the center of colonies that displayed low H3K9me3 tended to be bunched closely together and spatially corresponded to sites of apical actin rings with converging stress fibers (Figure 5B, S4A, S4B, S4C). To confirm this, we measured total H3K9me3 levels in early differentiating colonies categorized based on their actin cytoskeleton morphology as either ‘apical constriction’ or ‘relaxed’ (Figure 5B). This showed that compared to naïve colonies, relaxed early differentiation colonies had reduced levels of H3K9me3, while colonies with features of apical constriction showed the even lower H3K9me3, indicating that apical constriction during early differentiation correlates with loss of a repressive histone modification. We next sought to determine if loss of H3K9me3 during apical constriction was mediated by contractility and LINC-dependent mechanotransduction. We found that inhibition of contractility with blebbistatin or LINC perturbation with DN-SRKASH expression abolished the reduction in H3K9me3 that occurred in constricted, early differentiating colonies compared to naïve colonies (Figure 5C). Thus, apical constriction morphogenesis during exit from naïve pluripotency drives loss of the repressive histone modification H3K9me3 via contractility- and LINC-mediated mechanotransduction.

**Figure 5.**
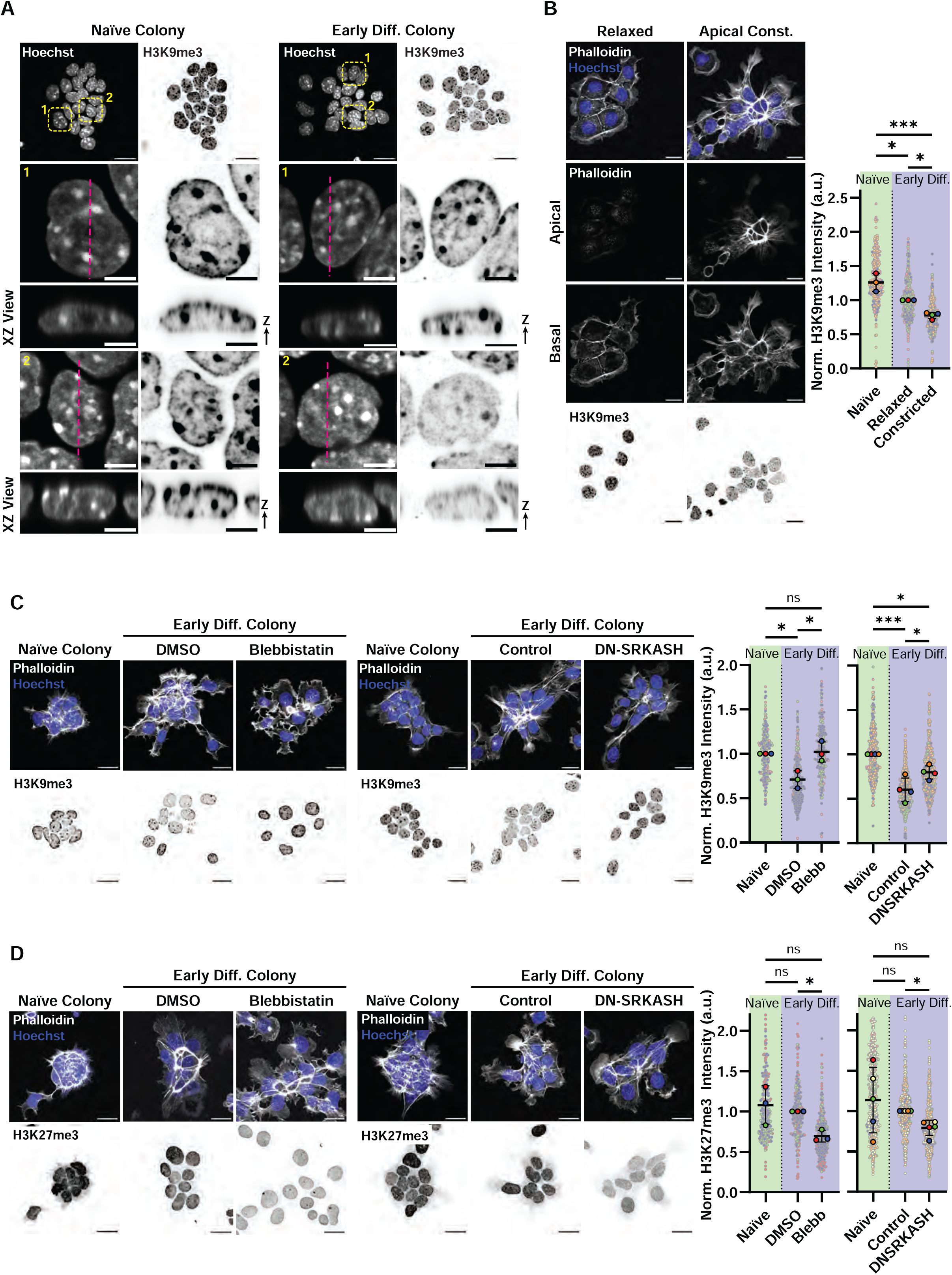
Apical constriction morphogenesis drives changes in H3K9me3 and H3K27me3 histone modifications in a contractility- and LINC-dependent manner. **(A-D)** Confocal images (**B-D**, left) of mESCs fixed in media containing inhibitors (2i) to maintain naïve pluripotency (Naïve Colony, **A**, **C, D**) or at 16 hours after 2i withdrawal to allow exit from naïve pluripotency and initiation of early differentiation (Early Diff. Colony), immunostained for H3K9me3 (**A-C**, inverted grayscale) or H3K27me3 (**D**, inverted grayscale) and stained with Hoechst to visualize DNA (**A**, grayscale, **B-D**, blue) and fluorescent phalloidin to label F-actin (**B-D**, grayscale), Scale bars, 20μm, or 5μm for zoomed single nuclei. **(A)** Top row: 3D projection of super-resolution confocal stack, dashed yellow boxes highlight regions zoomed (2^nd^ and 4^th^ rows)and pink dashed lines highlight the plane of the XZ view of the 3D images shown as XZ views (3^rd^ and 5^th^ rows) . **(B)** Left panels: examples of cell clusters lacking (Relaxed, left column) or bearing (Apical Const., right column) actin features of apical constriction; Top row: 3D projection of confocal stack. Middle row: apical and Bottom row: basal confocal slices. Right: Quantification of normalized (relative to relaxed) mean of max intensity Z projections of individual nuclei (mean ± SD, N=3 independent experiments, colored points correspond to individual cells within each experiment, Repeated Measures Mixed effects analysis, Tukey Multiple Comparisons Test, *P=0.0109, **P=0.0002, *P=0.0158). **(C, D)** Cells were treated with either 2μM blebbistatin or DMSO vehicle control (left panels) or lacking (control) or bearing inducible RFP-tagged DN-SRKASH with doxycycline treatment (middle panels). Right: Quantification of normalized mean fluorescence intensity mean of maximum intensity Z projections of individual nuclei for **(C)** H3K9me3 (relative to Naïve control) or **(D)** H3K27me3 (relative to Early Diff DMSO or Control lacking DN-SRKASH). (**C,** mean ± SD, N=3 and N=4 independent experiments, Repeated Measures Mixed effects analysis, Tukey Multiple Comparisons Test, Blebbistatin: *P=0.0488, *P=0.0392, DN-SRKASH: ***P=0.0008, *P=0.0219, *P=0.0263) **(D,** mean ± SD, N=3 and N=5 independent experiments, paired two-tailed t test, Blebbistatin: *P=0.0324, DN-SRKASH: *P= 0.0167

Loss of the strongly repressive H3K9me3 on histones can sometimes be compensated by an enrichment of H3K27me3, another repressive modification that may poise genes for subsequent expression, and this has notably been observed in pluripotent stem cells and epidermal progenitor cells (Fukuda et al., 2023; Le et al., 2016; Neagu et al., 2020; Peters et al., 2003; Zhang et al., 2022). We thus sought to determine if apical constriction-mediated mechanotransduction could drive the replacement of repressive H3K9me3 with H3K27me3 to poise genes for later expression. Immunostaining for H3K27me3 in naive cells along with visualization of F-actin and DNA showed that this histone modification was generally dispersed in the nucleoplasm, with a subset of cells displaying enrichment at the nuclear periphery (Figure 5D, S4D, S4E, S4F). While induction of exit from naïve pluripotency caused no significant change in H3K27me3 signal compared to naïve colonies, inhibition of either contractility using blebbistatin or LINC perturbation with DN-SRKASH expression during early differentiation resulted in reduction of H3K27me3 compared to controls (Figure 5D). Super-resolution imaging showed that nuclei expressing DN-SRKASH gained H3K9me3 and lost H3K27me3 compared to controls (Figure S4G, S4H). We conclude that morphogenesis-associated contractility during early differentiation drives loss of H3K9me3 while it maintains high levels of H3K27me3, and these activities are mediated by the LINC complex, which may have implications on lineage commitment via regulation of developmental gene expression.

### The *Sox1* Locus Translocates To The Nuclear Periphery During Early Differentiation, Where Actomyosin Contractility And The LINC Drive H3k27me3 Enrichment On Its Promotor

Given our finding that neurectoderm lineage commitment driven by SOX1 expression depends on apical constriction-mediated nuclear mechanotransduction, we hypothesized that forces on the nucleus could physically drive movement of the *Sox1* locus to position it for subsequent activation. To test this, we employed 3D DNA fluorescence in situ hybridization (FISH) and 3D confocal imaging (Figure 6A, B) in cells immunostained for lamin B1 to mark the position of the NE. This showed that in the naïve condition, although inactive, both alleles of the *Sox1* locus were positioned in the nuclear interior at a similar distance from the NE as both alleles of the active *Pou5f1* (OCT4) control gene (Figure 6C). In contrast, during early differentiation, both alleles of the *Sox1* locus were significantly closer to the NE than they were either in naïve cells or compared to the position of the Pou5f1 loci at the same timepoint. Later in differentiation, around the time of Sox1 transcriptional activation (72hr), the *Sox1* loci returned to a more central position, similar to that of the *Pou5f1* loci. We also tested whether contractility or the LINC complex played a role in *Sox1* locus positioning but found that neither blebbistatin treatment nor DN-SRKASH expression caused significant changes in either *Sox1* or *Pou5f1* loci positions (Figure S5A). Thus, we conclude that during exit from naïve pluripotency when apical constriction is occurring, the *Sox1* locus is translocated to the nuclear periphery by a contractility- and LINC-independent mechanism.

**Figure 6.**
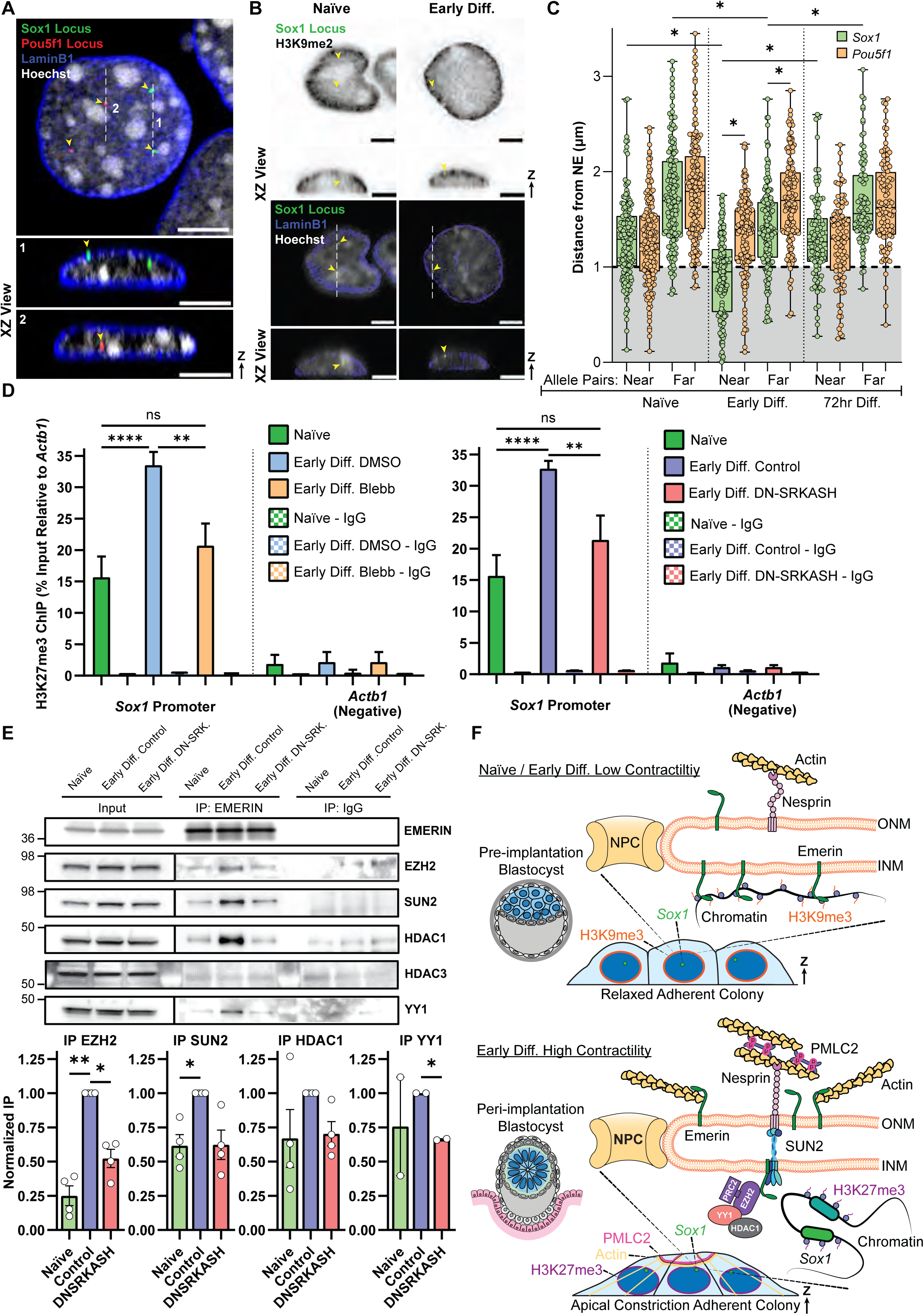
The *Sox1* locus resides at the nuclear periphery during exit from naïve pluripotency and requires actomyosin contractility and the LINC complex for H3K27me3 enrichment. **(A, B)** Super-resolution **(A)** and standard resolution **(B)** confocal images of mESCs fixed in media containing inhibitors (2i) to maintain naïve pluripotency (Naïve, **B**) or at 16 hours after 2i withdrawal to allow exit from naïve pluripotency and initiation of early differentiation (Early Diff.) immunostained for LaminB1 (blue **A, B**) and H3K9me2 (inverted grayscale, **B**) and stained with Hoechst to visualize DNA (grayscale) with genomic loci for *Pou5f1* (OCT4) (red, **A**) and *Sox1* (green) localized with 3D DNA FISH (yellow arrowheads point to FISH-labelled genomic loci, with 2 alleles visible per gene). White dashed lines highlight the plane of the XZ view of the 3D images below. Scale bars, 5μm. **(C)** Quantification of the distance of *Pou5f1* (OCT4) and *Sox1* genomic loci center to the nearest NE were measured in mESCs labeled as above in media containing 2i inhibitors (naïve), during early differentiation (16 hours after 2i withdrawal) and at 72 hours after 2i withdrawal to induce differentiation. The ‘near’ and ‘far’ allele measurements within the same nucleus were recorded separately. (Boxplots of the 25-75 percentile range while whiskers show total data range, N=3 independent experiments, Kruskal-Wallis test with Dunn’s multiple comparison Test, *P<0.0005). **(D)** Quantification of ChIP-qPCR from H3K27me3 or IgG control immunoprecipitations for sequences within the *Sox1* or *Actb1* promoters in naïve or early differentiating cells (16 hours after 2i removal, Early Diff.) in the presence of either 2μM blebbistatin or DMSO vehicle control (left) or of cells lacking (control) or bearing inducible RFP-tagged DN-SRKASH with doxycycline treatment (right) (mean ± SD of N=3 independent experiments, Ordinary one-way ANOVA with Tukey multiple comparisons test, Blebbistatin: \*\*\*\**P*<0.0001, \*\**P*=0.0021, DN-SRKASH: \*\*\*\**P*<0.0001, \*\**P*=0.0052). **(E)** Above: Western blot of immunoprecipitations with anti-emerin or equivalent anti-IgG antibodies from mESCs either before (Naïve) or 16 hours after 2i withdrawal (Early Diff.) to allow exit from naïve pluripotency and initiation of early differentiation of cells lacking (control) or bearing inducible RFP-tagged DN-SRKASH (DN-SRK.) with doxycycline treatment, probed with antibodies to EZH2, SUN2, HDAC1, HDAC3, and YY1. Below: Quantification of blots like those described above (mean ± SD, N=3 independent experiments, points correspond to mean each experiment, Repeated Measures One-Way ANOVA, Tukey Multiple Comparisons Test, EZH2: **P=0.0039, *P=0.0117, SUN2: *P=0.0351, YY1: *P=0.0211). **(F)** Hypothetical model of how apical constriction during early differentiation after exit from naïve pluripotency regulates *Sox1* for later expression to drive lineage commitment. See discussion for description.

Our observation of apical constriction-driven changes in histone modifications and contemporaneous movement of the *Sox1* locus suggested that the licensing of *Sox1* for later expression might be through the regulation of H3K27me3 enrichment on the gene promoter. Indeed, this enrichment has been previously shown to be required for subsequent activation of *Sox1* in pluripotent cells (Cruz-Molina et al., 2017). To test this, we performed ChIP-qPCR of H3K27me3 to examine its abundance in the vicinity of the *Sox1* promoter. This analysis showed an increase of H3K27me3 occupancy during early differentiation compared to the naïve condition (Figure 6D, S5B) (Kinoshita et al., 2021). This increase of H3K27me3 occupancy on the *Sox1* promoter was mitigated when contractility was inhibited with blebbistatin or when the LINC complex was disrupted with DN-SRKASH. These results indicate that the enrichment of H3K27me3 on the *Sox1* promoter during early differentiation depends on nuclear mechanotransduction.

The known interaction of emerin with the histone-modifying PRC2 complex (Demmerle et al., 2012, 2013; Haque et al., 2010; Ma et al., 2019; Marano & Holaska, 2022) together with our findings of emerin-dependent regulation of lineage commitment suggested the hypothesis that nuclear mechanotransduction may regulate the emerin-PRC2 interaction to effect chromatin modifications to mediate SOX1 expression in early differentiation. To test this, we analyzed the effects of early differentiation and LINC complex perturbation on known emerin interacting partners by immunoprecipitation (Figure 6E). This analysis suggested that compared to the naïve state, during early differentiation emerin increased interactions with the LINC component, SUN2, the PRC2 component, EZH2, and the PRC2-recruiting enzyme with DNA sequence specific binding protein, YY1 (Caretti et al., 2005; Verheul et al., 2020; Wilkinson et al., 2006; Yi et al., 2021; Zhang et al., 2016). Although emerin has been reported to interact with HDAC3 (Demmerle et al., 2012), we found that HDAC3 was not detected in the emerin pulldown, however HDAC1 was present (Liu et al., 2015), and its abundance appeared to increase in a LINC-dependent manner in early differentiating compared to naïve cells. Force imparted on the LINC complex has been previously shown to promote emerin phosphorylation (Guilluy et al., 2014), however probing the emerin immunoprecipitates with phospho-specific antibodies failed to show differentiation- or mechanotransduction-dependent changes in tyrosine, threonine or serine phosphorylation of emerin (Figure S5C). These results suggest that during early differentiation, emerin engages with the SUN2 component of the LINC complex and recruits DNA binding and histone modifying enzymes, while a subset translocates to the ONM. Overall, our results suggest that during early differentiation, gene loci for the neurectoderm lineage factor *Sox1* translocate to the NE while apical constriction morphogenesis activates a contractility-, LINC-, and emerin-dependent nuclear mechanotransduction pathway that recruits histone modifying enzymes to poise *Sox1* for later expression during lineage commitment.

## Discussion

In this study, we tested the hypothesis that forces driving morphogenesis in pluripotent stem cells are transmitted to the nucleus and spur changes in nuclear architecture that are critical for pluripotency maturation and initial lineage commitment. During implantation, pluripotent cells of the epiblast undergo a maturation process to gain lineage competence, during which they rearrange from a cell mass to a strongly constricted apical surface, forming a rosette structure which gives rise to an epithelial layer (Bedzhov & Zernicka-Goetz, 2014; Neagu et al., 2020; Shahbazi et al., 2017). Although the genetic, molecular epigenetic, and morphological changes that occur during these early stages of embryogenesis are well-described, whether they are interdependent is not known (Marks et al., 2012; Smith, 2017; Yang et al., 2019). We utilized a chemically defined *in vitro* mESC model that can be triggered to undergo changes in gene expression that mirror the early differentiation of pluripotent cells of the peri-implantation epiblast *in utero* (Mulas et al., 2019; Ying et al., 2008). We show that triggering early differentiation of mESCs causes them to recapitulate the morphodynamic features of those in the peri-implantation epiblast, mirroring rosette formation, apical constriction and epithelialization. In adherent mESCs, this drastic cytoskeletal remodeling is marked by formation of apical, F-actin rich contractile purse-string structures and stress fibers that overlie and directly deform nuclei, along with the formation of perinuclear actin networks that enshroud the nuclear surface. By perturbing actomyosin contractility and known mediators of nuclear mechanotransduction, we discovered that apical constriction morphogenesis plays no role in exit from naïve pluripotency itself, but instead licenses an essential lineage commitment factor for later expression. Indeed, our studies revealed that there is a time window during early differentiation, coincident with apical constriction morphogenesis, during which *Sox1* is licensed by contractility-, LINC complex-, and emerin-mediated nuclear mechanotransduction for expression later in development to allow neuroectodermal lineage commitment. Our observation that over-expression of Lamin A, a component that is typically very low in pluripotent cells, also impaired later SOX1 expression, additionally implicates composition of the nuclear lamina being critical for *Sox1* regulation (Constantinescu et al., 2006; Eckersley-Maslin et al., 2013). 3D DNA FISH and analysis of nuclear architecture revealed that during apical constriction in early differentiation, the *Sox1* locus is translocated to the nuclear periphery and nuclear mechanotransduction drives a widespread loss of the constitutive heterochromatin mark, H3K9me3, and a compensatory accumulation of the facultative histone modification, H3K27me3, which was also enriched specifically on the *Sox1* promoter, licensing it for later expression (Cruz-Molina et al., 2017). Protein interaction studies further uncovered emerin-mediated engagement with the LINC complex and recruitment of DNA binding and histone modifying enzymes, including the H3K27me3-catalyzing EZH2, likely driving the local priming of the *Sox1* locus situated near the NE for later expression and lineage commitment (Boyer et al., 2006; O’Carroll et al., 2001; Wiles & Selker, 2017). Our study thus uncovers a potential role of tissue morphogenesis-driven nuclear mechanotransduction pathways that enable regulation of lineage gene expression, carrying implications for developmental processes that involve distinct cell shape changes with subsequent fate choices.

Based on the findings of our study and the work of others, we propose a hypothetical model where during early pluripotent cell differentiation, cell intrinsic morphogenic forces initiate a nuclear mechanotransduction cascade that is required for subsequent neuroectodermal lineage commitment (Figure 6F). First, in the naïve state, mESCs exhibit low contractility and nuclei maintain H3K9me3 through the enzymatic activity of Suv39h1/h2 and Setdb1, which function to maintain pluripotency gene expression and stabilize repetitive DNA (Karimi et al., 2011; Nicetto et al., 2019; Rea et al., 2000; Zhang et al., 2024). Upon Wnt signal withdrawal, morphogenesis of mESCs during early differentiation is initialized by cell polarization through PAR3/PAR6-mediated cell-cell adhesions and maintained via laminin ECM engagement of integrin β1, regulated by small GTPase Rap1, with subsequent apical constriction morphogenesis that shapes polarized cells into a rosette, followed by lumenogenesis driven by OTX2 expression (Bedzhov & Zernicka-Goetz, 2014; Kim et al., 2022; Lin et al., 2019; Mole et al., 2021; Neagu et al., 2020; Shahbazi et al., 2017). Apical constriction morphogenesis likely occurs via Cdc42/MRCK kinase-mediated specific targeting and activation of non-muscle myosin II to apical F-actin rich purse-string structures that delineate the converging apical surfaces of individual cells (Chu et al., 2013; Lee et al., 2007; Martin & Goldstein, 2014; Nakajima & Tanoue, 2010, 2011; Nishimura & Takeichi, 2008; Perez-Vale et al., 2023). Apical constriction morphogenesis generates forces within and between cells that are transmitted to the nucleus via F-actin induced deformations of the nucleus and the LINC complex (Crisp et al., 2006; Hoffman et al., 2020; Padmakumar et al., 2005). This force transmission to the nucleus results in a widespread loss of H3K9me3 in heterochromatin, including at the nuclear periphery, which is likely mediated by Piezo1-mediated calcium released from endoplasmic reticulum stores, and the rapid regulation of histone methyltransferases or demethylases (Nava et al., 2020; Song et al., 2022; Xu et al., 2023). Force transmission to the nucleus also stabilizes the SUN2 LINC components in the NE, and regulates emerin via a phosphorylation independent mechanism, instead possibly through modulating its oligomeric state, resulting in its enrichment in the outer nuclear membrane (Essawy et al., 2019; Fernandez et al., 2022; Guilluy et al., 2014; Le et al., 2016; Niu et al., 2022). Enrichment of ONM emerin could occur via sliding across membrane adjacent to nuclear pore complexes and could be stabilized via an increased capacity for its binding to the pointed end of actin filaments, which may allow free-barbed ends to grow and assemble an F-actin shroud that encases the nucleus and possibly amplifies nuclear mechanosensing and protects the nucleus from morphogenesis-associated damage (Holaska et al., 2004; Le et al., 2016; Ohba et al., 2004; Shao et al., 2015; Wales et al., 2016). The resulting, diminished emerin population retained in the INM gains capacity for increased interactions with SUN2, HDAC1 and H3K27me3-catalyzing EZH2 by an as yet unknown mechanism (Caretti et al., 2005; Liu et al., 2015; Ma et al., 2023; Marano & Holaska, 2022; Wang et al., 2011). In a process independent of myosin contractility and the LINC complex and tied to pluripotency progression, the locus of the master neuroectodermal lineage transcription factor, *Sox1,* moves to near the NE during early differentiation. Trapping of the *Sox1* locus at the nuclear periphery could be mediated by the DNA motif targeting factor that recruits lamin-associated sequences to the nuclear periphery, YY1, whose interaction with emerin we find may be driven by morphogenesis-associated nuclear mechanosensing (Donohoe et al., 1999; Harr et al., 2015). While situated at the nuclear periphery, *Sox1* is subject to mechanically induced changes in chromatin architecture via emerin recruitment of HDAC1 and EZH2. This results in enrichment of H3K27me3 on the *Sox1* promoter region, a prerequisite for its later expression (Cruz-Molina et al., 2017). There is also a more widespread enrichment of H3K27me3, including the periphery, to maintain heterochromatin repression to compensate for the mechanically induced, widespread loss of H3K9me3 (Neagu et al., 2020; Pailles et al., 2022; Peters et al., 2003; Saksouk, Barth, et al., 2015). However, H3K27me3 enrichment on the *Sox1* promotor functions to maintain its repression and prevent premature activation until a later stage in development, where the appropriate neural lineage cues, such as loss of BMP and Wnt signals, gain of FGF signals, and the active E2A transcription factor, become present (Rao et al., 2020; Shparberg et al., 2019). These cues would otherwise fail without the permissive regulatory topology maintained by H3K27me3 on the *Sox1* promoter (Cruz-Molina et al., 2017).

Overall, we present findings that demonstrate a coordination between tissue morphogenesis in the very earliest stages of development and the regulation of lineage gene expression, mediated by cell-intrinsic actomyosin forces imparted on the nucleus via the LINC complex. Throughout development, as cells generate forces to change shape and thereby determine higher order tissue architecture, our findings implicate the potential role of forces transmitted to the nucleus in spurring chromatin changes downstream of NE mechanotransduction to drive subsequent gene expression choices. Elucidating mechanotransduction-driven fate choices in other contexts of developmental morphogenesis would help determine generality of such mechanisms in development, and thus allow the control of stem cell differentiation, thereby contributing to understanding of congenital diseases and genetic conditions where morphogenesis or nuclear components are affected.

## Acknowledgements and Declaration of Interests

We thank the Waterman Lab for project support, and members, especially William Shin, Dr. Ana Pasapera, Dr. Robert Fischer, the National Heart, Lung, and Blood Institute (NHLBI), and the NHLBI light microscopy core, especially Dr. Xufeng Wu. We thank Dr. Xuefei Ma for the kind gift of their NM-MyoIIB antibody. We additionally thank the Cambridge Stem Cell Institute and Core Facilities, especially microscopy support from Peter Humphreys, and members of the Basu Lab and Chalut Lab, especially Dr. Carla Mulas and Dr. Lawrence E. Bates. This work was supported by the Division of Intramural Research Program at the NHLBI (CMW), and an ERC Consolidator Grant (772798, KJC). SB was part-funded by the UKRI Biotechnology and Biological Sciences Research Council (BBSRC) (BB/W000423/1), the Trinity College Stem-Cell Medicine Senior Postdoctoral Researcher Fund and a starting grant at the Cambridge Stem Cell Institute from the Wellcome Trust (203151/Z/16/Z) and UKRI Medical Research Council (MC_PC_17230). DY was supported by Wellcome Trust PhD programme in Stem Cell Biology and Medicine (218481/Z/19/Z). GWW was supported by the EPSRC studentship. YAM was supported by the Max Planck Society. MSH was supported by the NHLBI and the NIH Oxford Cambridge Scholars Program. Declaration of interests: KJC is a co-founder and CSO of Cyclana Bio, GWW is Technical Director of Thomas Keating Ltd, and remaining authors declare no competing interests.

## Materials and Methods

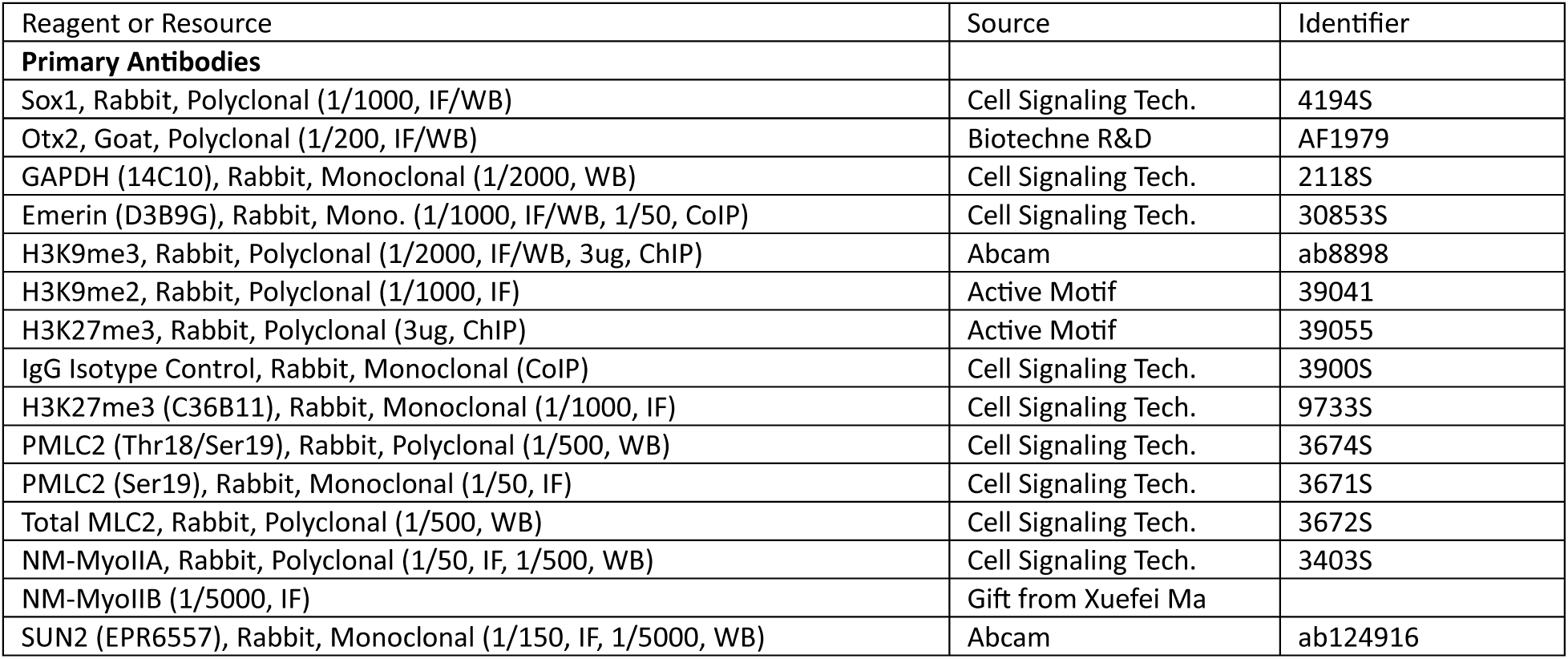
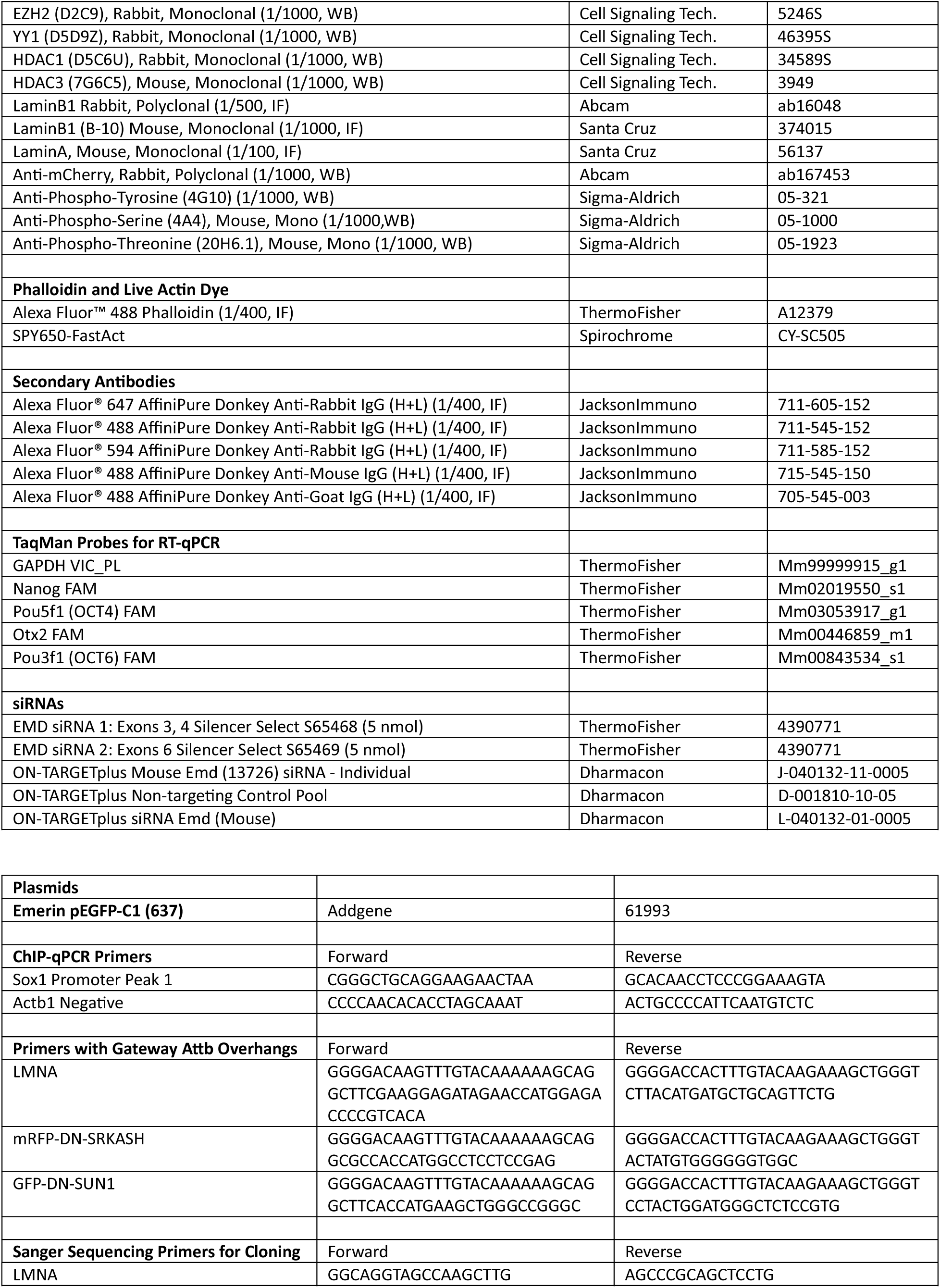
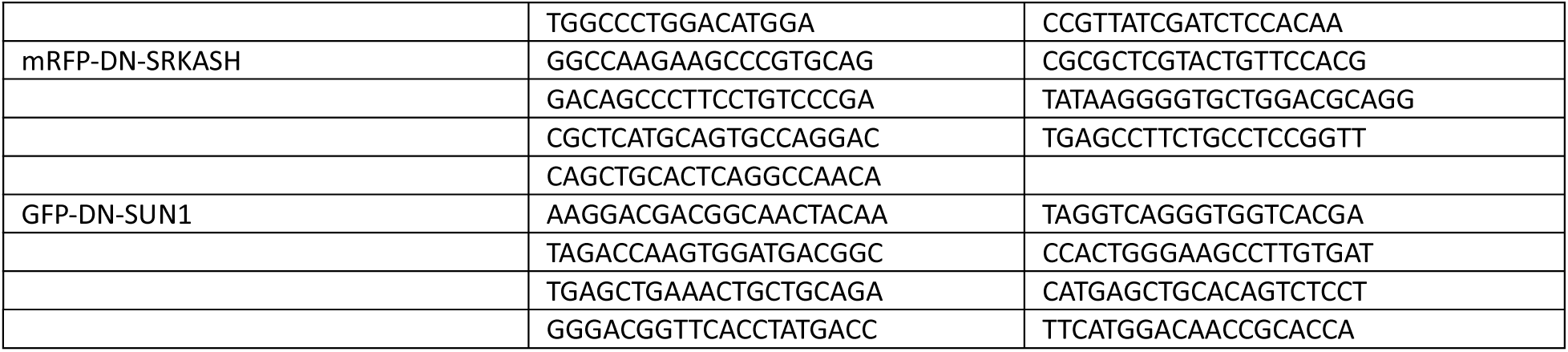
Key Resources Table

### Mouse Embryonic Stem Cell Culture

E14Tg2a (E14) (Hooper et al., 1987) mESCs were previously derived from embryos and stored in liquid nitrogen in the Cambridge Stem Cell Institute in “2i+Lif” culture media to maintain naïve pluripotency. For routine maintenance prior to experiments, cells were plated on 0.1% Gelatin-coated (Sigma-Aldrich G1890), tissue culture-treated plastic culture dishes at a density of 15,000 to 20,000 cells per square centimeter. 2i+Lif culture media was made in house with a base of “N2B27” media supplemented with MEK/ERK inhibitor PD0325901 (1uM, Axon Medchem 1408), GSK3β inhibitor CHIR99021 (3uM, Axon Medchem 1386), and mouse LIF (10 ng/mL, Recombinant mouse Leukemia Inhibitory Factor protein, Qkine Qk018). The base N2B27 was made from equal parts DMEM/F-12 (ThermoFisher 21041025) and Neurobasal media (ThermoFisher 12348017), along with Gibco B27 (0.5x, ThermoFisher 17504044), human insulin (12.5ug/mL, ThermoFisher 12585014), L-glutamine (0.78mM, or 2mM final with DMEM, ThermoFisher 25-030-081), Beta-Mercaptoethanol (100uM, ThermoFisher 21985023), and homemade N2 supplement, which contained Apo-transferrin human (8.76mg/mL, Sigma-Aldrich T1147), Sodium Selenite (0.54ug/mL, Sigma-Aldrich S5261), Putrescine (1.68mg/mL Sigma-Aldrich P5780), Progesterone (2.05ug/mL Sigma-Aldrich P8783), and 0.657% BSA Fraction V (0.657% ThermoFisher 15260037) diluted in DMEM/F-12. N2B27/2i+Lif media formulation and in house formulation for N2 (BV.N2) was derived from the following publication: (Mulas et al., 2019).

Cells were routinely split every 2-3 days. Cell colonies were dissociated into single cells by pipetting in Accutase (ThermoFisher A1110501). Cells were pelleted with 300g centrifugation for 3.5 minutes. As the dynamics of early exit from naïve pluripotency was of interest, care was taken to prevent colony overgrowth, and colony roundness and presence of a refractile edge was used as an indicator for colony quality. Colony overgrowth, or poor media quality, was quickly indicated by unusual cell death or loss of refractile edges and roundness of colonies. In the case of overgrowth or poor media quality, the culture was discarded, and new cells were thawed and cultured in newly prepared media.

Cryopreservation of cells was achieved after Accutase dissociation of cells, and resuspension of cell pellet in 2i+Lif media with 10% DMSO as a cryoprotectant. 1mL of cell suspension was transferred into appropriate cryo-vials and cooled in an isopropyl alcohol containing chamber to slow the rate of cooling to -1°C per minute overnight at -80°C. The next day, cells were transferred into liquid nitrogen storage.

Prior to differentiation experiments, mESCs cultured in 2i+Lif media were instead cultured in 2i media without LIF for an additional passage. Addition of LIF delays the exit from naïve pluripotency by approximately 12 hours when compared to mESCs cultured in 2i without Lif (Mulas et al., 2019). For experiments studying the exit from naïve pluripotency and neuroectodermal differentiation, cells were replated at 10,000 cells per square centimeter (unless otherwise stated) on tissue culture treated plastic, or Poly-L-ornithine (0.01%, Sigma-Aldrich P4957)-treated glass for 1 hour at RT, and next coated with Laminin (10 ug/mL, Sigma-Aldrich Laminin from Engelbreth-Holm-Swarm murine sarcoma basement membrane L2020) in PBS (ThermoFisher 14190144), overnight at 4°C. This protocol was derived from protocols of monolayer neuroectodermal differentiations of mESCs from the following publications (Mulas et al., 2019; Ying et al., 2003). Upon replating, the cells are cultured in the base N2B27 media without the 2i inhibitors.

### Cloning and Generation of Inducible Expression Cell Lines

Cell lines for inducible overexpression of DN-SRKASH or LMNA are described in (Wylde, 2017). DNA constructs for the SRKASH and SUN1 dominant negative perturbation cell lines were kind gifts from Dr. Gant Luxton (UC Davis).

The DN-SRKASH consists of the coding sequence of nesprin-2G, but lacks the actin-binding calponin homology domain, and acts as a dominant negative which effectively decouples the LINC complex by outcompeting binding to endogenous SUN proteins (Lombardi et al., 2011; Stewart-Hutchinson et al., 2008). The cell line for inducible expression of DN-SUN1 was generated using a truncation of EGFP-SUN1, designated SS-EGFP-SUN1-457-913, which encodes amino acids 457-913 of SUN1 that lacks its transmembrane domain but contains the luminal KASH-binding SUN domain and has incorporated Signal Sequence for ER and inter-NE-space targeting (Hennen et al., 2018).

Expression plasmids were constructed with Invitrogen Gateway Technology with Clonase II (ThermoFisher 12535029). cDNAs encoding RFP- tagged DN-SRKASH, GFP- tagged DN-SUN1, and untagged LaminA were amplified from source plasmids, with PCR primers that included attB1 and attB2 overhangs for downstream Gateway reaction compatibility (sequences listed in Key Resources Table, under Primers with Gateway Attb Overhangs). PCR amplification of attB overhang products was achieved with Q5 Hotstart HiFi Polymerase (New England Biolabs M0493), following manufacturer’s protocol. Next, with attB overhang PCR products, a Gateway BP Clonase II (ThermoFisher 11789020) reaction was performed to insert genes into entry vector backbone, Gateway pDONR221 (ThermoFisher 12536017). Next, with the BP Clonase II reaction-derived entry vectors containing the originally amplified genes of interest, they were inserted into a destination vector 144 PB TetO (PB-TAP IRI)(Dos Santos et al., 2014) comprised of a doxycycline-inducible gene expression system with hygromycin resistance, also containing transposon elements compatible with PiggyBac transposase. Insertion of entry vectors into this destination vector was achieved with a Gateway LR Clonase II (ThermoFisher 11791100) reaction. Cells were transfected with the resulting expression vectors, along with 203 CAG rtTA3 with Blasticidin resistance with Invitrogen Lipofectamine 3000 (ThermoFisher L3000001) and appropriate antibiotic selection (150 µg/ml Hygromycin B and 10µg/ml Blasticidin). Cells were reverse transfected (Amarzguioui, 2004; Ovcharenko et al., 2005), where 40,000 Accutase-singularized cells were replated in 12 well plate wells in 500µL of media including 1.5µL of Lipofectamine 3000 reagent, 300ng of destination vector plasmid, 300ng of 203 CAG rtTA3 and 100ng of PBase (PiggyBac Transposase). The 203 CAG rtTA3 constructs encode for expression of the rtTA transcription factor, which is a required component of doxycycline-induced Tet-On system, which is constitutively expressed, and upon binding to doxycycline, will bind *tetO* sequences and initiate transcription. After antibiotic selection, surviving colonies with stable piggyBac transposase-mediated genome integration of the expression plasmid and rtTA were harvested and selected based on morphology and flow cytometry of EGFP or mRFP expression. For the untagged Lamin A line, multiple surviving clones were immunostained for Lamin A for selection.

Between cloning steps, amplicon and plasmid lengths were checked to verify successful PCR amplification and recombination reactions with 1% agarose gel electrophoresis with restriction enzyme digestions, using PflMI (New England Biolabs R0509) and Xhol (New England Biolabs R0146) in NEBuffer 3 (New England Biolabs B7003). All constructs generated were verified with Sanger sequencing, with sequencing primers listed in Key Resources Table. Plasmid propagation was achieved with DH5α competent *E. coli* (ThermoFisher 18265017), or One Shot ccdB Survival 2 T1R Competent Cells (ThermoFisher A10460). Plasmids were purified using Qiagen Plasmid Plus MIDI kit (Qiagen 12943), following manufacturers protocol.

### Proliferation rates

To measure the impact of blebbistatin treatment on cell proliferation rate, cells cultured in 2i with various concentrations of blebbistatin or DMSO vehicle were plated at 10,000 cells per square centimeter into 6 wells of a 12 well plate per condition. Cells were harvested in 20 hours intervals and counted while suspended in 1mL of Accutase. Cell counts were performed with a Beckman Coulter Vi-Cell XR Cell Viability Analyzer. (Beckman Coulter).

### Confocal and Live Cell Imaging

Spinning disk confocal imaging was conducted with a Nikon Eclipse Ti2 microscope with a Yokogawa CSU-W1 spinning disk scan head, paired with a Nikon Perfect Focus system and a C13440-20CU ORCA-Flash4.0 V3 Digital CMOS Camera. Illumination was provided by a Nikon LUNV 6-line laser unit (20 mW 405nm; 20 mW 445nm; 70 mW 488nm; 40 mW 515nm; 70 mW 561nm and 40 mW 640nm; power measurement is at the fiber output) with 2 single mode optical fiber outputs (1 APC and 1 UPC on a fast actuator). For the objective, a Nikon Plan Apo 60XA/1.40 Oil DIC H inf/0.17 WD 0.21 was used. For wider field of view confocal imaging to compare neighboring colonies, a Nikon Plan fluor 40x/1.30 Oil DIC H objective was used. This microscope was also used for epifluorescence imaging of SOX1 immunofluorescence using the Nikon Plan Fluor 10x/0.30 Ph1 DL Inf/1.2 WD 15.2 objective for wide field of view. The microscope was equipped with a Nikon motorized stage with XY linear encoders and a Mad City (Madison, WI) Nano-Z100 piezo insert with 200µm travel. Laser confocal illumination was selected with electronic shutters and an automated filter turret containing a multi-bandpass dichromatic mirror together with an electronic emission filter-wheel. The microscope was piloted with NIS-Elements software (Nikon). Z-stack images were taken with 0.3µm steps, and side profile images were generated with Fiji Java-based image processing software (Schindelin et al., 2012).

Live cell imaging was done with an identical microscope setup, except with a Nikon Eclipse Ti body equipped with a Yokogawa CSU-X1 spinning disk scan head and illuminated with an Agilent MLC400B Monolithic Laser Combiner with four lasers (405nm: 20mW, 488nm: 50 or 80mW, 561nm: 50 or 80mW, 647nm: 125mW) and a Photometrics CoolSNAP Myo CCD camera. This microscope was equipped with a Tokai Hit enclosed stage-top incubator system, calibrated to 37°C, humidified, and maintained a 5% CO2 atmosphere. Live cells were imaged on 35 mm glass bottom dishes (Cellvis D35-10-1.5-N).

Super-resolution imaging was done with an Airyscan-equipped Zeiss LSM880 and the Axio Observer7 confocal microscope stand equipped with a Wienecke & Sinske GmrH piezo-Z stage and piloted with ZenBlack software (Zeiss). The objective used was a Plan-Apo 63x 1.4 NA DIC M27 oil objective and the microscope was equipped with a 32-channel GaAsP-PMT area detector. Airyscan image reconstructions were processed in auto-strength mode using ZenBlack software (Version 2.3). 488 nm multi-line 25 mW Argon laser, 561 DPss solid state 15mW laser and 633nm HeNe 5mW lasers were used.

### Phalloidin and Immunofluorescence Imaging

For standard immunofluorescence, cells were fixed at room temperature for 10 minutes in 4% paraformaldehyde (PFA) in PBS, then washed three times in PBS containing 0.1% Tween-20 (PBS-T). Cells were then blocked for 1 hour at room temperature on a rocker in blocking buffer (5% Normal Donkey Serum, 0.3% TritonX-100 in PBS). Primary antibody dilution was then added in antibody dilution buffer (1% Bovine Serum Albumin, 0.1% TritonX-100 in PBS) and left overnight at 4°C on a rocker. The next day, samples were washed with PBS-T three times, then diluted secondary antibody was added and incubated in the dark for 1 hour on a rocker at room temperature. The sample was washed with PBS-T then stained with 20µM Hoechst 33342 (ThermoFisher 62249) diluted in PBS-T for 20 minutes at room temperature, then finally washed in PBS-T and kept in PBS for imaging.

Preservation of the f-actin cytoskeleton in mESCs required cytoskeleton-optimized buffer and fixation conditions. 10x concentrated PIPES Cytoskeletal Buffer (pCB) was prepared, containing 100mM PIPES pH6.8, 1380mM KCl, 30mM MgCl2, 20mM EGTA in MilliQ sterile water and filter sterilized. Cells cultured on “Squeaky-clean” glass coverslips (Waterman-Storer, 2001) were fixed with 4% PFA from a fresh ampoule diluted in 1x pCB that was pre-warmed in a water bath to 37°C and incubated at 37°C for 20 minutes. Next, cells were permeabilized with 0.25% Triton X-100 in pCB at room temperature for 10 minutes. Then, free aldehydes were quenched with a wash in in 0.5 mg/mL Sodium Borohydride in pCB at room temperature for 10 minutes. Cells were then washed twice with pCB and stored in 4°C in TBS-T (TBS with 0.1% Tween20) until further processing.

For F-actin staining, fixed cells were blocked in blocking buffer (0.05% Tween-20, 0.1% TritonX-100, 2% BSA, 2% Normal Donkey Serum, in 1x TBS) also containing a 1/400 dilution of Fluor 488 Phalloidin (ThermoFisher A12379) and incubated for 60 minutes at room temperature. Coverslips were then carefully placed cell face down on 100µL drops of diluted primary antibody in antibody dilution buffer (0.1% TritonX, 1% BSA in 1x TBS). They were kept in a humidified and sealed chamber and incubated overnight in 4°C. The next day, coverslips were washed in TBS-T three times for 5 minutes. Cells were then incubated in diluted secondary antibodies for 60 minutes in room temperature, then washed again three times, and finally incubated in 20µM Hoechst for DNA staining, washed a final time in TBS then mounted on microscope slides in Invitrogen SlowFade Gold Antifade Mountant (ThermoFisher S36936) and sealed with nail polish.

For selective immunostaining of emerin in the ONM, cells were subjected to identical immunofluorescence protocols except with the removal of tween-20 and triton-X to minimize permeabilization of the nuclear membrane. Instead, the permeabilization step was done with 40µg/mL digitonin diluted in PBS at 4°C for 4 minutes and washed with ice cold PBS.

A different cytoskeletal buffer and fixation protocol was followed to better preserve perinuclear f-actin structures, derived from the following publications (Le et al., 2016; Vinzenz et al., 2012). This protocol did not preserve epitopes as well as the PIPES-based cytoskeleton buffer and PFA fixation, hence was not usable for parallel immunofluorescence staining. 10x concentrated MES Cytoskeletal Buffer (mCB) was prepared, containing 100mM MES pH6.1, 1500mM NaCl, 50mM MgCl2, 50mM EGTA, 50mM Glucose in MilliQ sterile water and filter sterilized. For fixation, an initial fixation was done with 0.25% glutaraldehyde, along with 0.5% Triton X-100 in 1x mCB for 1 minute with the fixative prewarmed to 37°C. Following this, a second fixation was done with 2% glutaraldehyde with 1.2676 µM of Alexa Fluor 488 Phalloidin (ThermoFisher A12379) in 1x mCB, prewarmed to 37°C, and incubated at 37°C for 15 minutes. Cells were then washed in 0.5 mg/mL Sodium Borohydride wash in mCB at room temperature for 10 minutes, and finally incubated in 20µM Hoechst for nuclear staining, washed a final time in TBS then finally mounted on microscope slides in Invitrogen SlowFade Gold Antifade Mountant (ThermoFisher S36936) and sealed with nail polish.

### Image Analysis

In experiments determining ratio of SOX1 positive cells after 72 hours of 2i withdrawal and differentiation, cells for each condition were cultured in triplicate wells. In each technical replicate well, 10 epifluorescence images were taken with only the Hoechst channel viewed. Images were segmented into individual nuclei with CellProfiler image analysis software (Stirling et al., 2021) from Hoechst-labelled nuclei channel images, with a three class Otsu intensity-based thresholding followed by intensity based clumped object segmentation. With individual nuclei segmented from the Hoechst channel, the average intensity of SOX1 was measured in the corresponding immunostaining channel on a per nucleus basis. Cells in the naïve 2i condition were immunostained for SOX1 and imaged with same settings to determine background levels of SOX1 signal, which was used as the cutoff for positive SOX1 expression.

To measure the intensity of perinuclear actin, mid colony-height optical sections from spinning-disk confocal Z-stacks were used. A 4-pixel wide area was drawn around the outer perimeter of the Hoechst-labelled nuclei channel. Areas of nuclei perimeters that overlapped with other bright actin structures, such as lamellipodia, cell-cell junctions or stress fibers, were not traced to ensure the measurement was solely of distinct perinuclear actin networks. The average intensity of the phalloidin channel was determined, and measurements were normalized to the naïve condition.

To determine total levels of H3K9me3, SUN2, and emerin, spinning disk confocal z-stacks were taken of colonies using the phalloidin channel. Maximum projections of 3D images were produced with Fiji, then individual nuclei were segmented from the Hoechst image channel with CellProfiler with a two class Otsu intensity-based thresholding followed by shape-based clumped object segmentation. CellProfiler was then used to measure the mean intensity of SUN2 and corresponding mRFP-DNSRKASH or emerin and corresponding laminB1, or H3K9me3 fluorescence intensities for every segmented nucleus.

In all plots comparing differences of immunofluorescence intensity, data was represented in the style of “SuperPlots”, where statistics were performed on the means of repeated experiments, displayed as prominent dots, with lighter shaded dots of corresponding color in the background representing the measurements of individual cells (Lord et al., 2020).

### Western Blot Analysis

Cells were plated in 10cm dishes, and proteins were extracted and scraped on ice using 50µL of SDS Sample Buffer (2x) with fresh 2-Mercaptoethanol (5%). Samples were then sonicated with a probe sonicator on ice in a cold room (4°C) three times for 5 second intervals and boiled at 95°C for 5 minutes. Samples were loaded onto a Tris-Glycine 4-20% gradient gel (ThermoFisher XP04200BOX) with SeeBlue Plus2 Prestained Protein Standard (Thermofisher LC5925). Gels ran for 90 minutes at 100 volts. After, gels were transferred to PVDF (Thermofisher 88518) membranes with wet transfer at 15 volts for 2 hours. Membranes were blocked in TBS with 0.1% Tween-20 (TBS-T) with 5% milk (or 5% BSA for phosphorylated proteins except PMLC) for one hour at room temperature on a rocking table before adding primary antibodies at their proper dilution in 5% BSA overnight at 4°C on a rocking table. Treated membranes were then rinsed, then washed three times in TBS-T on a shaker table for 5 minutes each wash. Species-appropriate HRP-conjugated secondary antibodies in 5% BSA were then added and incubated for one hour in room temperature on a rocking table. Membranes were again rinsed and washed three times, then treated with Immobilon Western Chemiluminescent HRP Substrate (Sigma-Aldrich WBKLS0500) before imaging on an iBright FL1500 Imaging System. Quantification of western blots was performed from digital images using Fiji software with its Gel Analyzer tool with local background subtraction adjacent to bands of interest.

Genomic loci distance measurements were measured in 3D using Bitplane IMARIS software from 3D reconstructions of confocal image stacks. With IMARIS, the distance of the center of each genomic locus was measured to the nearest nuclear periphery. The nuclear lamina surface was generated using the Surfaces tool with automatic settings based on the fluorescent signal from LaminB1 immunofluorescence channel, with the surface filled with automatic settings based on the Hoechst channel, while the 3D DNA FISH genomic loci were localized using the Spots tool with a 300nm diameter centered at peak fluorescence intensity. The distance from the FISH loci center (both alleles for each gene) to the nearest LaminB1 surface was measured with the Shortest Distance to Surfaces tool.

### RNA isolation and relative quantitative real-time PCR

Cell lysates were captured directly in Qiagen RLT Buffer Plus from the RNA Plus Mini Kit (Qiagen 74134) and RNA was isolated according to manufacturer’s protocol. Reverse transcription reactions were carried out with SuperScript IV Reverse Transcriptase (Thermofisher 18090010), containing 1µL of 10mM dNTP mix (Thermofisher R0192), 1uL of 50µM Random Hexamers (Thermofisher N8080127), added to 100ng of purified RNA and annealed at 65°C for 5 minutes. Next, 1µL of 100mM DTT, 1µL of RNaseOUT (Thermofisher 10777019), and 1µL of enzyme in 1x SSIV Buffer were added to the reaction, with a final volume of 20µL, which was set to 55°C for 10 minutes and 80°C for 10 minutes to carry out reverse transcription to generate cDNA and enzyme inactivation. Next, relative quantitative real-time PCR was performed in a total reaction volume of 10µL, using 2µL cDNA, 0.5 µL gene-specific TaqMan FAM probe, 0.5uL of GAPDH TaqMan VIC probe, 5uL TaqMan Fast Advanced Master Mix (Thermofisher 4444556) and 2µL of water (TaqMan probes listed in Key Resources Table). Reactions were done in technical triplicate, with the quantification of gene expression acquired with a Roche LightCycler 96 real-time PCR instrument, with a thermal reaction profile of 50°C for 2 minutes, 95°C for 20 seconds, and 40 cycles of 95°C for 1 second followed by 60°C for 20 seconds. Gene expression values were expressed as a fold-change relative to RNA from control (equivalent vehicle control, non-treated cells), normalized against GAPDH using the ΔΔCT method (Livak & Schmittgen, 2001).

### 3D#DNA FISH

OligoPaint probes were designed using the PaintShop toolkit (https://paintshop.io/) to obtain a set of probes targeting 50kb regions around *Sox1* and *Pou5f1* (OCT4) gene start sites (chr8:12385771-12436732 and chr17:35483404-35533404, respectively). Oligonucleotide probe sets of ∼250 probes for each gene with at least 10bp of spacing between subsequent probes were obtained from Twist Bioscience (Beliveau et al., 2012; Nguyen & Joyce, 2019). Probe amplification was achieved with T7 amplification as previously described (Beliveau et al., 2017).

Probe hybridization was performed on cells fixed and stained for LaminB1 or H3K9me2 with our standard immunofluorescence protocol, followed by a post-fix with 2% PFA for 10 minutes at room temperature (RT). Cells were permeabilized again with 0.7% Triton-X for 10 min at RT, followed by 5 min at RT incubations, first in 2x SSC buffer with 0.2% Tween-20 (SSCT), followed by 2x SSCT with 50% formamide. Chromatin was denatured by incubating cells in 2x SSCT with 50% formamide for 2.5 minutes at 92C, then at 60C for 20min. Then, after briefly cooling the sample to RT, a hybridization mix, containing 50% formamide, 10% dextran sulfate, 10ug RNaseA, 5mM dNTPs, ∼100-200pmol of gene targeting probes, and 6pmol of signal-amplifying bridge oligos was added and sealed with rubber cement, and incubated for 2.5min at 92C on a heat block, then the sample was allowed to hybridize with the probe overnight at 37C. The next day, the cell samples were washed with 2x SSCT for 15min at 60C, then 10 min in 2x SSCT at RT, then again in 0.2x SSC at RT then finally in 2x SSC. A secondary probe hybridization mix, containing 10% formamide, 10% dextran sulfate, and 45pmol of AF488 and AF647 fluorescently labelled hybridizing oligos was applied to the cells, and incubated in a humidified chamber for 2 hours at RT. Cells were finally washed for 5min in 2x SSCT at 60C, then again at RT, then stained for Hoechst and mounted on glass slides for imaging.

### ChIP-qPCR

A minimum of 10 million cells for each condition were collected from 15cm culture dishes. Cells were fixed for 4 minutes at room temperature on a shaker table with 1% PFA from fresh ampoule (Electron Microscopy Sciences 15710) diluted in PBS. Fixation solution was then replaced with a stop fixation buffer (0.125M Glycine in PBS) and incubated for 5 minutes at room temperature. Stop fixation solution was then aspirated, and cells were rinsed twice with ice cold PBS. Next, cells were scraped in ice cold scraping solution (1x Protease Inhibitor Cocktail Roche 11697498001 and 1mM PMSF in PBS). Scraped cells were kept on ice followed by centrifugation to pellet cells in a swinging bucket rotor at 550g for 15 minutes at 4°C. Supernatant was carefully discarded, and 1µL of 100x PIC and 100mM PMSF was added to the pellet, then it was snap frozen in dry ice and stored in -80°C.

For processing, thawed pellets were resuspended in ice cold lysis buffer (25mM Hepes pH7.9, 1.5mM MgC2, 10mM KCl, 0.1% NP-40 in MilliQ Sterile Water) and incubated for 40 minutes on ice with occasional shaking. Nuclei were released by homogenization of the lysed cell suspension with 10 strokes on ice in a Dounce B homogenizer. The homogenized, lysed cell suspension was then transferred to a new 1.5mL tube and centrifuged for 4600g for 10 minutes at 4°C. The supernatant was aspirated, and the nucleus pellet was then resuspended in 130µL of sonication buffer (50mM Hepes pH7.9, 140mM NaCl, 1mM EDTA, 1% TritonX-100, 0.1% Na-Deoxycholate, 0.1% SDS in MilliQ water). The sample was then loaded into 130µL wells of Covaris Sonication Tube Strip (8 microTUBE-130 AFA Fiber H Slit Strip V2 520239). The Covaris ME220 Focused-Ultrasonicator was used to shear samples down to DNA fragment sizes of 200-600BP with the following optimized settings (Temperature: 6-12°C, Base Pair Mode: 350bp, Repeat/Iterations: 6x, Repeat Process Treatment Duration: 10 seconds, Peak Power: 70 Watts, Duty Factor: 20%, Cycles Per Burst: 1000x, Total Treatment Time Per Sample: 60s). The sheared sample was then loaded into a new 1.5mL tube and spun for 21,000g for 10 minutes at 4°C, which left the sheared chromatin in the supernatant. 6µL was taken as an input sample that was purified for DNA to check for correct sheared DNA size with an Agilent 4200 TapeStation system with High Sensitivity D1000 ScreenTape (Agilent 5067-5584)and concentration with a Qubit DS High-sensitivity assay (ThermoFisher Q33230). The remaining chromatin was equally divided for immunoprecipitation experiments. For each condition, an equivalent amount of chromatin was taken for isotype IgG control, with equivalent amount of antibody mass added. Before antibody incubation, chromatin samples were topped up to 500µL with Sonication buffer with 1x PIC and 1mM PMSF. Sheared chromatin and antibody mixes were incubated overnight in a cold room rotating in an end-over mixer. Protein A Dynabeads (Thermofisher 10001D) were washed with Sonication Buffer two times and resuspended in Sonication Buffer. 30µL of beads were added to each sample, and they were again rotated for 4 hours in the cold room. Tubes were then placed on a magnetic rack for washing. They were washed once with sonication buffer, once with wash buffer A (50mM Hepes pH7.9, 500mM NaCl, 1mM EDTA, 1% Triton X-100, 0.1% Na-Deoxycholate, 0.1% SDS in MilliQ Sterile Water). They were then washed in Wash Buffer B (20mM Tris pH8.0, 1mM EDTA, 250mM LiCl, 0.5% NP-40, 0.5% Na Deoxycholate in MilliQ Sterile Water). Then washed in TE Buffer, then finally eluted in 200µL of Elution Buffer (TE Buffer with 1% SDS). Beads were incubated in elution buffer for 30 minutes at 65°C. Input and pulldown chromatin was finally purified into DNA with standard reverse crosslinking with Proteinase K (Thermofisher EO0491), and DNA purification with Phenol:Chloroform (Thermofisher 15593031) extraction.

Resulting purified DNA fragments were used for SYBR green based RT-qPCR with PowerTrack SYBR Green Master Mix (Thermofisher A46109). Reactions were performed in technical triplicate, with 10uL reactions containing 5ng cDNA, and 8uM of forward and reverse primers (primer sequences listed in Key Resources Table). A Roche LightCycler 96 real-time PCR instrument was used to quantify relative abundance of DNA fragments containing primer target sequences from anti-H3K27me3 immumoprecipitates, input control, and IgG pulldown control products, with a thermal reaction profile of 95°C for 2 minutes, and 40 cycles of 95°C for 15 seconds followed by 60°C for 1 minute. ChIP-qPCR results were analyzed as percentage of input by normalizing ChIP-qPCR to input DNA of the target gene, relative to *Actb1* as a negative control.

### Co-Immunoprecipitation

Immunoprecipitation of endogenous emerin was performed as follows. Cells were seeded in 15cm tissue culture dishes, and at least 15 million cells were collected for each condition. First, cells were rinsed with PBS containing 1x Protease Inhibitor Cocktail (PIC, ThermoFisher 11697498001), 1mM Phenylmethylsulfonyl fluoride (PMSF) and 1x phosphatase inhibitor cocktail (PhIC, ThermoFisher A32957). Next, the cells were scraped in ice cold PBS containing freshly added PIC, PMSF and PhIC, and pelleted at 550g for 15 minutes in 15mL falcon tubes in a swinging rotor bucket to minimize aggregation of cells on tube walls. With supernatant carefully aspirated, the pellet was snap-frozen in liquid nitrogen and stored at -80°C. Next, the pellet was thawed and resuspended in 500uL of ice-cold Lysis Buffer (50mM Tris-HCL pH7.4, 150mM NaCl, 1mM EDTA, 0.5% Triton X-100, 1mM DTT in MilliQ sterile water) containing freshly added PIC, PMSF and PhIC. This was incubated on ice for 25 minutes with occasional flicking. Samples were transferred to 1.5mL Eppendorf tubes and sonicated with probe sonicator on ice in the cold room three times for 5 second intervals. Protein concentrations were determined with a Pierce 660nm protein assay kit (ThermoFisher 22662) and volumes were adjusted to have equal protein concentrations and final volume of 525uL (at ∼450ug/mL). 25uL was taken as an input sample, and the rest was precleared with 30uL of washed Protein A Dynabeads (ThermoFisher 10001D) and rotated in an end-over mixer at 4C for 30 minutes. The sample was then evenly split for 0.1ug emerin antibody (CST Emerin (D3B9G) XP® Rabbit mAb #30853S) immunoprecipitation or equivalent mass of Rabbit IgG isotype control (Rabbit (DA1E) mAb IgG XP® Isotype Control #3900S). This was incubated overnight at 4°C on a spinning rotor. The next day, 40uL of washed Protein A Dynabead pellets were resuspended in the lysate and antibody immunocomplex mix and then incubated again on a spinning rotor for 4 hours at 4°C. Beads were then pelleted on a magnet and washed 5 times with 500uL of ice cold wash buffer (50mM Tris-HCL pH7.4, 150mM NaCl, 1mM EDTA, 0.05% Triton X-100 in MilliQ sterile water) on ice. Dynabead pellets were finally resuspended in 50uL of 2x SDS Sample Buffer containing 0.715M Beta-Mercaptoethanol to elute bead-bound proteins, and 50uL was also added to input samples, and samples were heated to 95-100°C for 5 minutes. Finally, the magnetic beads were pelleted, and the supernatant was transferred to new tubes and analyzed by western blotting as described above.

### siRNA Knockdown

siRNA knockdown of emerin was performed with Lipofectamine RNAiMAX reagent (Thermofisher 13778075) following steps in the manufacturer’s protocol. Optimizations were made to minimize cell death in mESCs. The lipofectamine was diluted 2000x. Final siRNA concentration was 60nM. To ensure sufficient protein knockdown for early exit from naïve pluripotency, mESCs were dissociated with Accutase into single cells, and were reverse transfected (Amarzguioui, 2004; Ovcharenko et al., 2005) in 2i media to maintain naïve pluripotency on laminin-coated dishes. Cells were left in transfection media overnight for 16 hours, then media was carefully replaced with fresh 2i media. To mitigate additional cell divisions from additional time in 2i culture, cells were plated at 8,000 per square centimeter. Six hours later, 2i media was removed and cells were washed with PBS and replaced with N2B27 to initiate exit from naïve pluripotency and neuroectodermal differentiation. siRNA knockdown efficiency was confirmed with immunofluorescence imaging or western blot analysis.

### Statistical Analysis

All statistical analysis was done using GraphPad Prism 10 software, with statistical tests and number of independent experiments listed in figure legends. Significance is reported in the figure legends with P>0.05 being not significant (N.S.).

**Figure S1.**
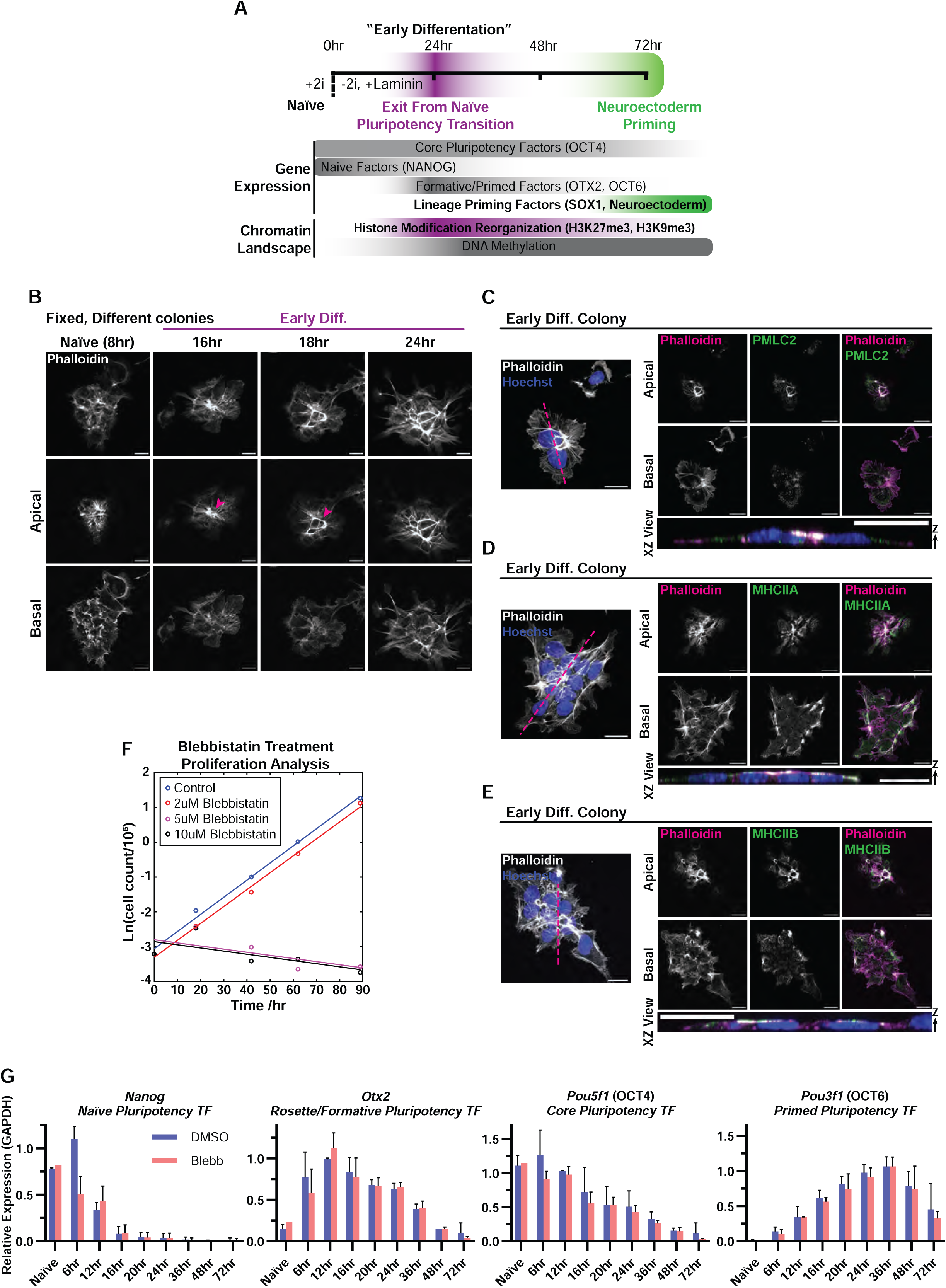
Apical Constriction morphogenesis is recapitulated by adherent mESCs during the exit from naïve pluripotency, and contractility is not required for pluripotency progression. (A) Schematic of the experimental design for investigating exit from naïve pluripotency in mESCs. mESCs maintain naïve pluripotency adhered to gelatin substrates in media containing inhibitors of MEK/ERK and GSK3β (2i, PD032590 ‘PD03’ and CHIR99021 also known as ‘Chiron’, respectively). When replated on laminin with 2i inhibitors withdrawn, mESCs begin to exit naïve pluripotency, a transition marked by the downregulation of the naïve factor NANOG, the maintenance of the core pluripotency factor OCT4, and the upregulation of the formative and primed pluripotency factors OTX2 and OCT6. This transition is also marked by global changes in chromatin, including a departure from a global DNA hypomethylated state, and a redistribution of repressive histone post translational modifications, H3K9me3 and H3K27me3. By default, mESCs exiting naïve pluripotency induced by 2i withdrawal commit to the neuroectodermal lineage, marked by the expression of master lineage transcription factor, SOX1. **(B)** Confocal images of adherent, mESC colonies fixed at the noted times after 2i withdrawal to allow exit from naïve pluripotency (Naïve) and initiation of early differentiation (Early Diff) with F-actin labelled with fluorescently-tagged phalloidin (greyscale). Top rows show 3D projections of confocal Z-stacks, second and third rows show apical and basal confocal images respectively. Pink arrowheads point to bright apical F-actin ring structures. Scale bars, 10μm. **(C, D, E)** mESC colonies fixed 16 hours after 2i withdrawal (Early Diff.) to allow exit from naïve pluripotency and initiation of early differentiation and stained with fluorescently-tagged phalloidin to visualize F-actin (grayscale or magenta) and Hoechst to visualize DNA (blue) with immunostaining of **(C)** PMLC2, **(D)** non-muscle myosin heavy chain isoform IIA (MHCIIA, grayscale or green) or **(E)** non-muscle myosin heavy chain isoform IIB (MHCIIB, grayscale or green). 3D projection of confocal stacks (left), apical confocal sections (upper row right), basal confocal section (middle rows right) and pink dashed lines highlight the plane of the XZ view of the 3D images (bottom row right). **(F)** Proliferation rates of mESCs with 2i media removed and replated at time zero, plotted on a log scale against time. mESCs were treated with different concentrations of Blebbistatin (2μm, 5μm and 10μm) at time zero, and compared to an untreated Control, which had a doubling time of 14.1 hours. Cells treated with 2μm had a doubling time of 14.3 hours. **(G)** RT-qPCR time course of mESCs in the presence of DMSO control (DMSO) or 2μM blebbistatin (Blebb, added at time 0 hours) at the noted times after 2i withdrawal to allow exit from naïve pluripotency (Naïve) and initiation of early differentiation, measuring RNA expression levels of naïve (Nanog), intermediate (Otx2), primed (Pou3f1 or OCT6) and core (Pou5f1 or OCT4) pluripotency factors (mean ± SD, N=3 independent experiments).

**Figure S2.**
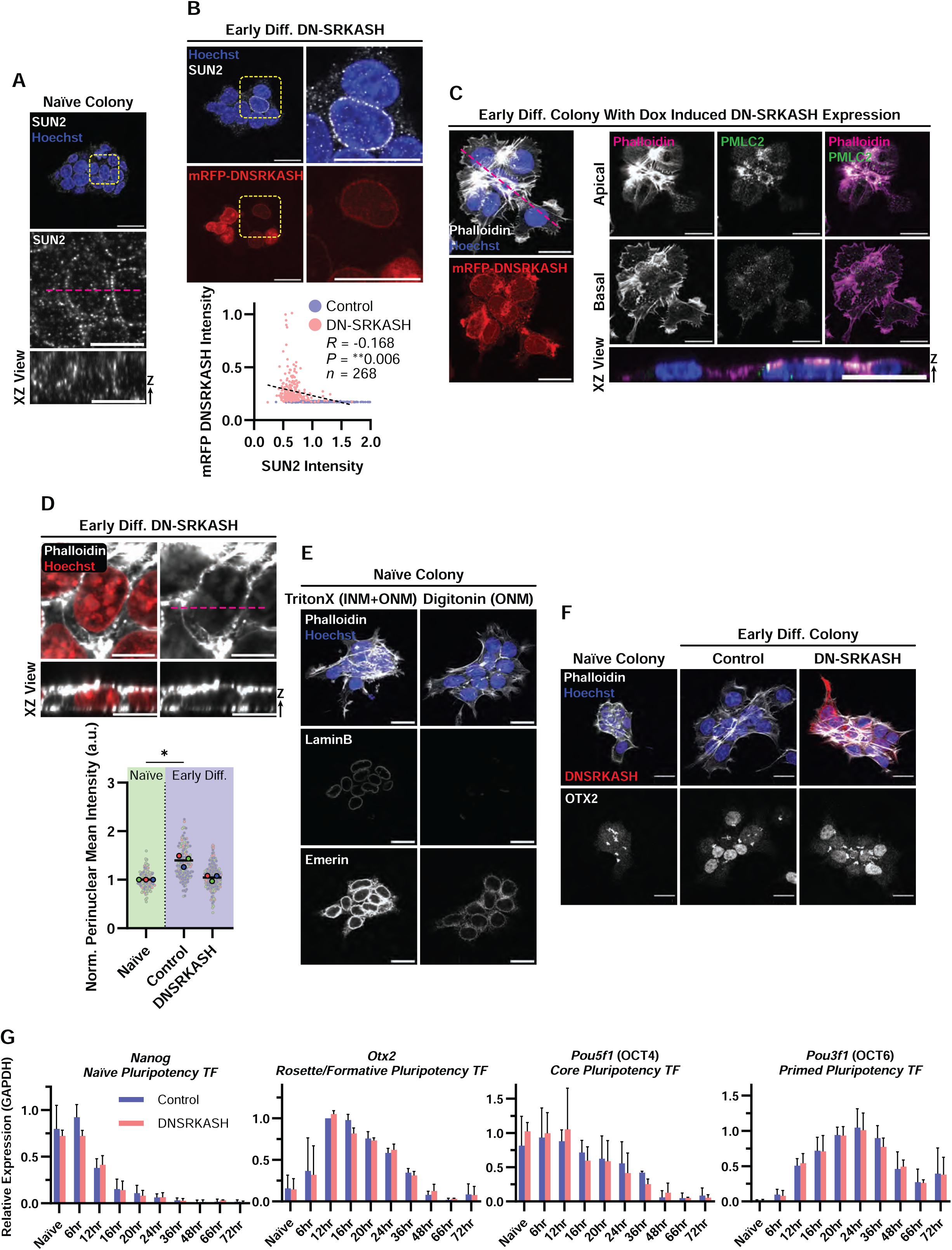
DN-SRKASH expression permits apical constriction morphogenesis and progression of pluripotency during early differentiation, while inhibiting SUN2 at the NE and perinuclear actin enrichment. **(A-F)** Confocal images of fixed mESC colonies either lacking (control) or bearing inducible RFP-tagged DN-SRKASH (red in **B, C, F**) with doxycycline treatment, fixed in 2i (Naïve) or after 16 hours of 2i withdrawal to allow exit from naïve pluripotency and initiation of early differentiation (Early Diff) and stained with fluorescently tagged phalloidin to visualize F-actin (grayscale or magenta) and Hoechst to visualize DNA (blue in **A, B, C, E, F**, red in **D**) with additional immunostaining of **(A, B)** SUN2 (grayscale, mid height colony optical sections), **(C)** PMLC2 (grayscale or green, left panels showing projections of 3D confocal Z-stacks, top and second rows show apical and basal confocal images respectively), **(E)** emerin & LaminB1 (grayscale, with top panels as projections and other panels mid colony height cross sections, Cells were permeabilized with tritonX-100 (left column) to allow antibody labelling of both the INM and ONM, or digitonin (right column) to allow antibody labelling of the ONM but not the INM or **(F)** OTX2 (bottom row, projections of 3D confocal Z-stacks) **(A, C, D)** Pink dashed lines highlight the plane of the XZ view of the 3D images in the bottom rows. **(A, B)** Yellow dashed box indicates region of zoomed nuclei below **(A)** or at right **(B)**. Scale bars, 20μm, while zoomed nuclei in **(A, B, D)** scale bars, 5μm. **(B)** Bottom panel: Correlation analysis between DN-SRKASH expression level and SUN2 intensity at NE. (n=319 cells for Control, n=268 for DN-SRKASH with Pearson correlation testing, Two-tailed, **P=0.0057) **(D)** Bottom panel: Plot of perinuclear actin intensity (mean ± SD of N=3 independent experiments, Ordinary one-way ANOVA with Tukey multiple comparisons test, \**P*=0.0488). **(G)** RT-qPCR time course of mESCs in the presence (DNSRKASH) or absence (control) of doxycycline to induce expression of DN-SRKASH at the noted times after 2i withdrawal to allow exit from naïve pluripotency (Naïve) and initiation of early differentiation, measuring RNA expression levels of naïve (Nanog), intermediate (Otx2), primed (Pou3f1 or OCT6) and core (Pou5f1 or OCT4) pluripotency factors. (mean ± SD, N=3 independent experiments).

**Figure S3.**
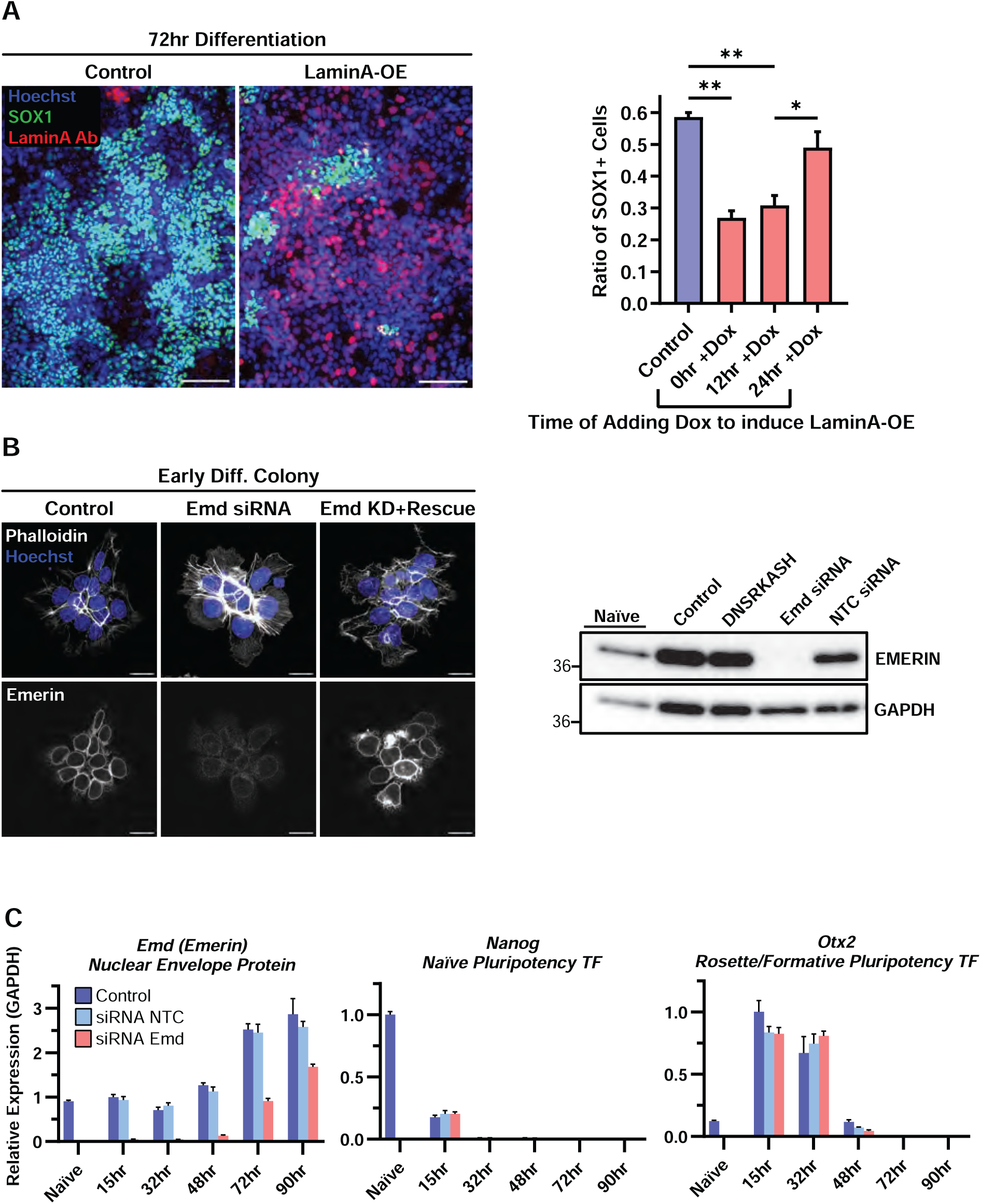
Ectopic overexpression of LaminA inhibits SOX1 expression during an early differentiation window and Emerin siRNA knockdown validation. **(A)** Left: Epifluorescence images of mESCs fixed after 72 hours that were cultured in media containing inhibitors (2i) to maintain naïve pluripotency (Naïve Condition) or after 2i withdrawal to allow exit from naïve pluripotency and early lineage commitment, immunostained for SOX1 (green) and LaminA (red) and stained with Hoechst to visualize DNA (blue) for cells lacking (control) or bearing inducible, ectopic expression of LaminA (LaminA-OE) with doxycycline (+Dox) treatment. Scale bars, 100μm. Right: Quantification of the fraction of SOX1-positive cells in samples fixed after 72 hours in 2i inhibitor (Naïve) or after 2i withdrawal to allow exit from naïve pluripotency and initiation of early differentiation. (mean ± SD of N=3 independent experiments, Ordinary one-way ANOVA with Tukey multiple comparisons test, **P<0.0021, **P=0.0034, *P=0.0164). **(B)** Left: Confocal images of mESC colonies fixed at 16 hours after 2i withdrawal to allow exit from naïve pluripotency and initiation of early differentiation (Early Diff.), mock-transfected (control), or transfected with siRNAs targeting the 3’UTR of emerin (*Emd* siRNA) alone or together with expression of emerin (Emd KD + Rescue) and stained with fluorescently tagged phalloidin to visualize F-actin (grayscale) and Hoechst to visualize DNA (blue), with emerin immunostained (bottom row). Top row shows 3D projections, while bottom row shows mid-colony height cross sections. Scale bars, 20μm. Right: western blot of lysates of mESC cultured in media containing inhibitors (2i) to maintain naïve pluripotency (Naïve) or mESC collected at 16 hours after 2i withdrawal to allow exit from naïve pluripotency and initiation of early differentiation (right 4 lanes), with above siRNA treatments or a non-targeting siRNA (NTC siRNA). Blots were probed for emerin and GAPDH as a loading control. **(C)** RT-qPCR time course data of mESCs exiting naïve pluripotency with above siRNA treatments, measuring RNA expression levels of naïve (Nanog), intermediate (*Otx2*) pluripotency factors, as well as *Emd* expression, in a 2i (naïve) state, or at the listed timepoints after 2i withdrawal.

**Figure S4.**
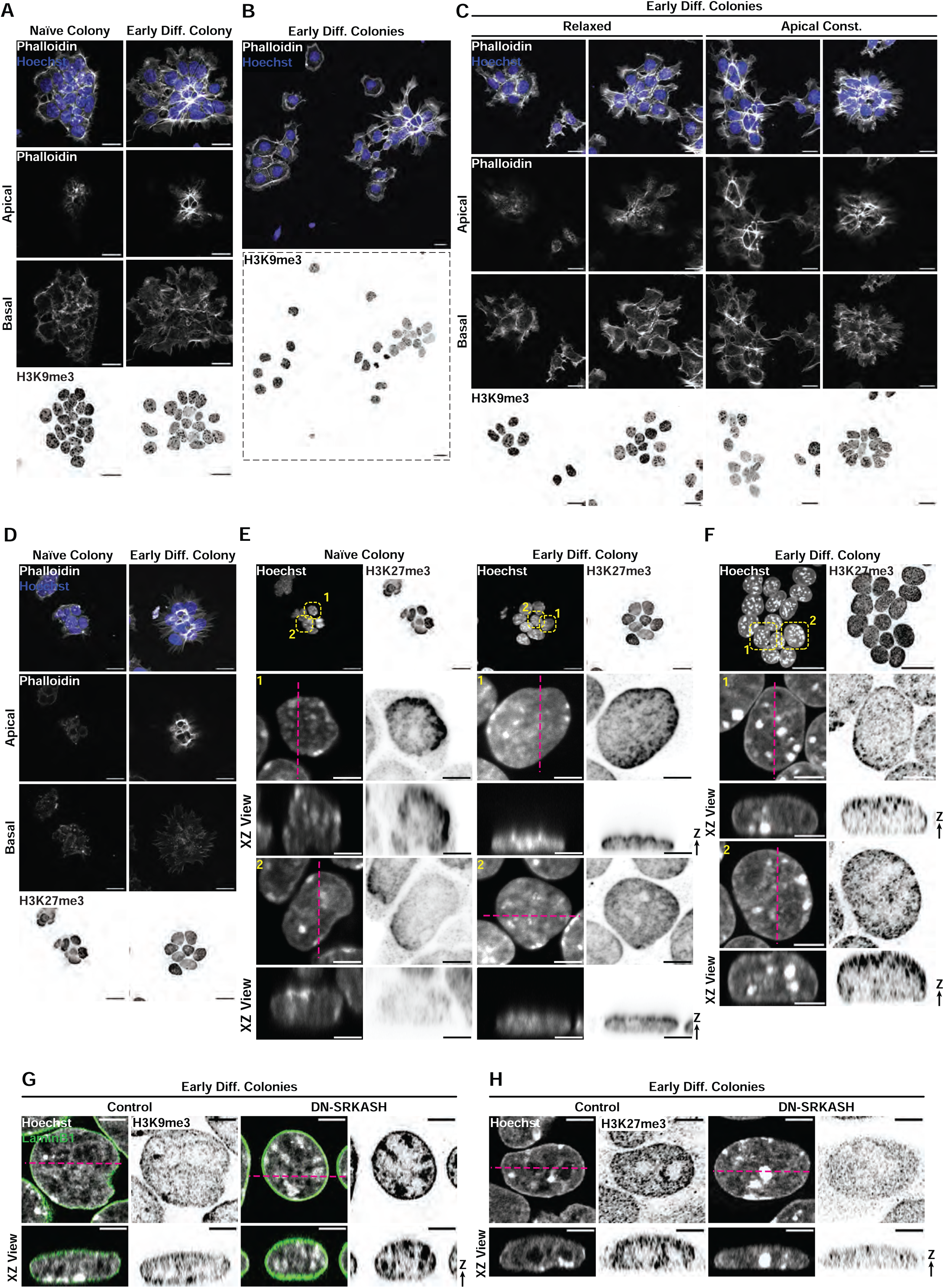
Apical constriction morphogenesis drives changes in histone modifications H3K9me3 and H3K27me3 in a contractility- and LINC-dependent manner. **(A-E)** Confocal or **(F-H)** super resolution confocal images of mESCs fixed in media containing inhibitors (2i) to maintain naïve pluripotency (Naïve, **A**, **D, E**) or at 16 hours after 2i withdrawal to allow exit from naïve pluripotency and initiation of early differentiation (Early Diff.), immu nostained for H3K9me3 (**A-C**, **G** inverted grayscale, projection) or H3K27me3 (**D**, **E, F**, **H**, inverted grayscale, projection) and stained with Hoechst to visualize DNA (**A-D**, blue, **E-H** grayscale) and fluorescent phalloidin to label F-actin (**A-D**, grayscale). **(A, C, D)** Top row: 3D projection of confocal stack, shown as XZ view of 3D image below. Second and third rows show apical and basal slices respectively. **(B)** Same example from Figure 5B, but with a wider field of view to show the cell clusters lacking or bearing actin features of apical constriction in the same field of view, projection of 3D images. **(E-H)** Top row: 3D projection of confocal stack, dashed yellow boxes highlight regions zoomed and pink dashed lines highlight the plane of the XZ view of the 3D images shown as XZ view of 3D image below. Scale bars, 20μm, or 5μm for zoomed single nuclei.

**Figure S5.**
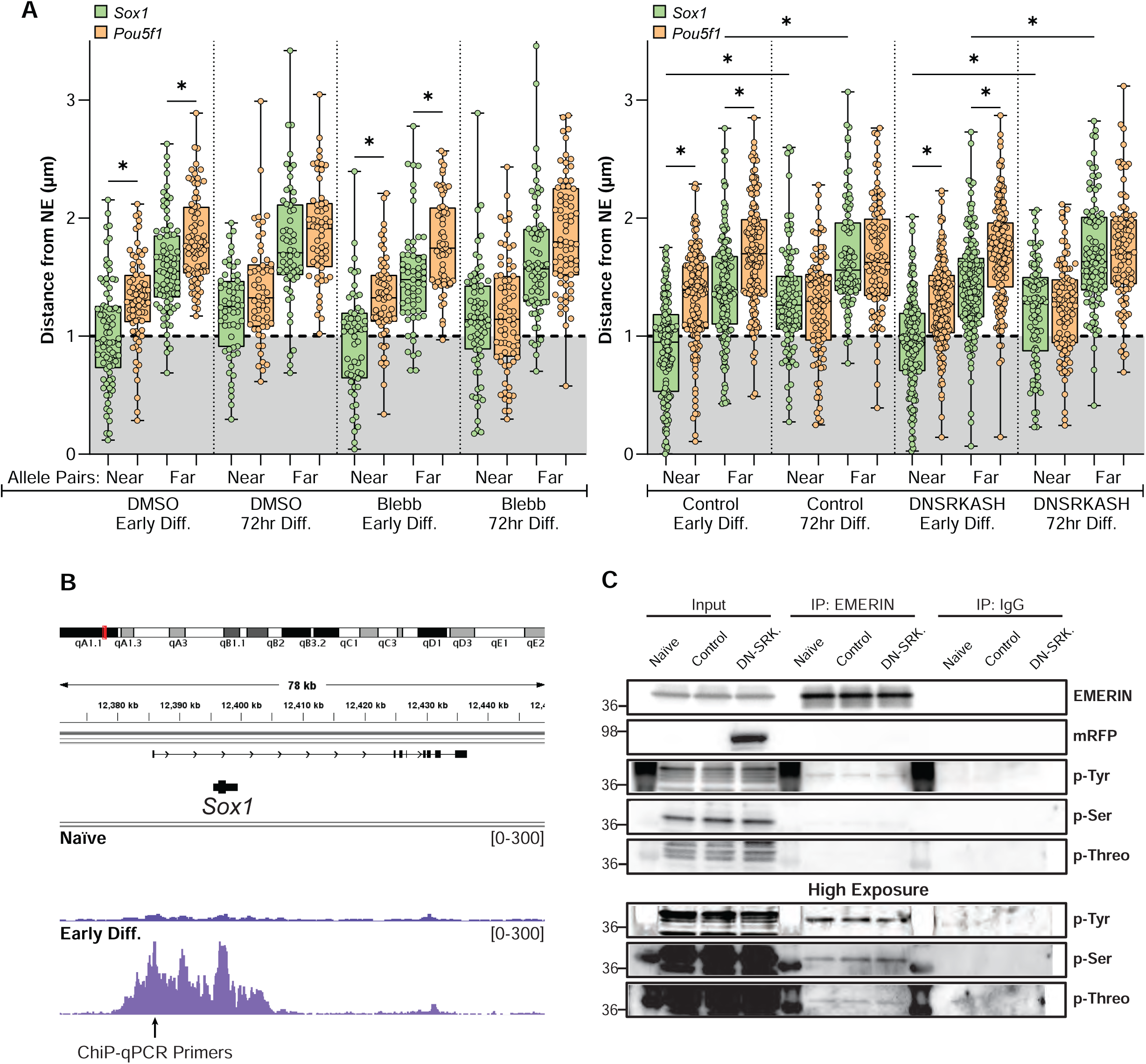
The *Sox1* locus resides at nuclear periphery during exit from naïve pluripotency, independent of contractility and LINC complex. **(A)** Quantification from 3D DNA fish experiments of cells immunolabeled for lamin B1 to mark the position of the NE of the distance of *Pou5f1* (OCT4) and *Sox1* genomic loci centers to the nearest NE were measured in mESCs labeled as in Figure 6A, B, in cells during early differentiation (Early Diff, 16 hours after 2i withdrawal) and at 72 hours after 2i withdrawal (72hr Diff.) to induce differentiation. The ‘near’ and ‘far’ allele measurements within the same nucleus were recorded separately. Measurements from mESCs in the presence of either 2μM blebbistatin (Blebb) or DMSO vehicle control (left) or of cells lacking (control) or bearing inducible RFP-tagged DN-SRKASH with doxycycline treatment (right, DNSRKASH) (Boxplots of the 25-75 percentile range while whiskers show total data range, Kruskal-Wallis test with Dunn’s multiple comparison Test, Blebb: *P<0.05, DNSRKASH: *P<0.005). (B) Chip-seq tracks from published datasets from (Kinoshita et al., 2021) showing levels H3K27me3 around the *Sox1* promoter, comparing mESCs in 2i (naïve) and after induction to differentiate towards a formative phase of pluripotency (Early Diff.). Red line indicates region on mouse chromosome 8 where *Sox1* resides, while the black arrow points to region probed by ChIP-qPCR primers for H3K27me3 enrichment at the *Sox1* promoter. Datasets visualized by Integrative Genomics Viewer (Robinson et al., 2011; Thorvaldsdottir et al., 2013). (C) Western blots of immunoprecipitations with anti-emerin or equivalent anti-IgG antibodies from mESCs either before (Naïve) or 16 hours after 2i withdrawal (Early Diff.) to allow exit from naïve pluripotency and initiation of early differentiation of cells lacking (control) or bearing inducible RFP-tagged DN-SRKASH with doxycycline treatment, probed with antibodies to phospho-Tyrosine, phospho-Serine, and phospho-Threonine. Bottom panels show a longer exposure of the same blot.

**Video S1.** Time-lapse (Confocal) of fluorescently labelled F-actin in live mESCs undergoing early differentiation upon replating and 2i withdrawal to induce exit from naïve pluripotency and early differentiation. Movie starts at 8 hours after 2i withdrawal. Images were captured every 10 min and ends after 30 hours (elapsed time shown in upper left). F-actin was labelled with Spirochrome SPY650-FastAct. The top left, top right, bottom left, and bottom right panels show a basal, medial, projection and apical slice of the same colony, respectively. Pink arrow highlights F-actin features of apical constriction. Scale bar, 20μm.

## Notes

### Summary of Updates

This version of the manuscript includes the reference list with no other changes.

